# Validating time-resolved fMRI with sub-second event durations

**DOI:** 10.1101/2023.07.30.550770

**Authors:** Alvin P.H. Wong, Esther X.W. Wu, Baxter P. Rogers, Christopher L. Asplund

**Affiliations:** Department of Psychology, Faculty of Arts and Social Sciences, National University of Singapore, 5 Arts Link, Singapore 117570, Singapore; N.1 Institute for Health and Institute for Digital Medicine (WisDM), National University of Singapore, 28 Medical Drive, #05-COR, Singapore 117456, Singapore; School of Psychology, Faculty of Health, Medicine and Behavioural Sciences, The University of Queensland, St Lucia QLD 4072, Australia; Division of Social Sciences, Yale-NUS College, National University of Singapore, 16 College Ave West, Singapore 138527, Singapore; Vanderbilt University Institute of Imaging Science, 1161 21^st^ Avenue South, AA-1105, Nashville, TN 37232, USA; Department of Radiology and Radiological Sciences, Vanderbilt University Medical Centre, 1211 Medical Center Drive, VUH 1145, Nashville, TN 37212, USA; Department of Biomedical Engineering, Vanderbilt University, PMB 351631, 2301 Vanderbilt Place, Nashville, TN 37235, USA; Department of Psychiatry and Behavioral Sciences, Vanderbilt University Medical Centre, 1211 Medical Center Drive, Nashville, TN 37232, USA; Department of Biomedical Engineering, College of Design and Engineering, National University of Singapore, 4 Engineering Drive 3, #04-08, Singapore 117583; Centre for Sleep and Cognition, Yong Loo Lin School of Medicine, 14 Medical Drive, #B1-01, Singapore 117599

**Keywords:** time-resolved fMRI, mental chronometry, peak latency, hemodynamic models

## Abstract

The hemodynamic signal measured with functional neuroimaging (fMRI) is notoriously sluggish, making it challenging to draw inferences about brief mental events. Relative timing differences in the response, however, can distinguish between changes in the duration or intensity of underlying neural activity (Henson et al., 2002). Specifically, increases in stimulus duration are predicted to delay the response peak and increase its amplitude, whereas increases in stimulus intensity should affect only peak magnitude (Friston, 2005). Although these different relationships have been empirically demonstrated using stimulus durations of several seconds, it remains unclear whether similar effects can be reliably detected on the sub-second timescales typical of cognitive events. Here we tested these predictions using brief visual and auditory stimuli that varied in intensity and duration. In Experiment 1 (n=15), stimuli were presented at three durations (100, 300, and 900 ms) and three intensities in a slow event-related design with a rapid TR (625 ms). In Experiment 2 (n=14), the stimulus durations were extended (1000, 1200, and 1800 ms) to enhance signal strength. Fitting the observed fMRI signals to parameterised hemodynamic response functions revealed that changes in stimulus duration affected both peak latency and magnitude, whereas changes in intensity affected only peak magnitude. Reliable effects were observed for duration differences as small as 200 ms, particularly when the evoked responses were strong. These findings support the validity of existing hemodynamic models and identify conditions under which sub-second timing differences can be reliably detected with fMRI. They also support the use of time-resolved fMRI as a principled and practical tool for mental chronometry in cognitive neuroscience.

## 1. Introduction

Using functional magnetic resonance imaging (fMRI) to characterize brief mental events presents a fundamental challenge: The BOLD (blood oxygen-level dependent) signal unfolds over several seconds, even when triggered by sub-second neural activity. Changes to either the intensity or duration of that activity are primarily reflected in the amplitude of the sluggish BOLD response. As such, BOLD amplitude is the key dependent measure in most neuroimaging studies, even when the manipulations are hypothesized to affect the duration of neurocognitive processes (Chen et al., 2023; Henson et al., 2002; Lindquist et al., 2009; Poldrack, 2015). Nevertheless, the BOLD signal contains information about the timing of mental events as well (Menon et al., 1998). Although duration and intensity changes have similar effects on the measured signal, relative timing differences can distinguish between these causes (Henson et al., 2002). Hemodynamic models and empirical findings show that an increase in stimulus duration leads to both an increase in the hemodynamic response’s peak latency (time to peak after the onset of the evoking stimulus) and peak magnitude, whereas increases in stimulus intensity lead to increased peak magnitude alone (Chen et al., 2023; Friston, 2005).

Many studies have relied on this dissociation between duration and intensity to test psychological theories. These theories typically involve bottlenecks of information processing, for which patterns of increased processing durations are the critical diagnostic feature. They therefore rely on the logic of mental chronometry, in which inferences about the properties of psychological events are based on changes in temporal measures, such as reaction time (Lo & Andrews, 2015). The hemodynamic response’s peak latency represents one such measure, with longer peak latencies thought to reflect increased processing durations. Peak latency shifts have been used to argue for, and to establish neural correlates of, bottlenecks in multitasking (Dux et al., 2006, 2009; Yue et al., 2025), attentional and response selection (Asplund et al., 2010; Marti et al., 2012; Scalf et al., 2011; Tombu et al., 2011), and language comprehension (Vagharchakian et al., 2012). In each case, the behavioral and peak latency timing differences across conditions were typically on the order of 200-1000 ms.

The magnitudes of these cognitively-relevant timing effects are considerably smaller than the empirical demonstrations linking even stimulus properties to hemodynamic response features. Distinctions between the duration and intensity of neural activity are reflected well in the BOLD signal with large timing differences (2 versus 6 seconds of contrast-reversing checkerboard stimuli, for example; Friston, 2005), but whether smaller differences yield reliably measurable effects is currently unknown. Prior relevant studies either used long stimulus durations or large stimulus duration differences (e.g. >1 s) (Liu & Gao, 2000; Vazquez & Noll, 1998), employed brief stimulus duration manipulations without varying stimulus intensity (Lewis et al., 2018), or did not compare effects of intensity and duration to determine whether they could be reliably distinguished (Vazquez & Noll, 1998; Yeşilyurt et al., 2008). The approach underlying the aforementioned bottleneck studies has not been empirically tested with the sub-second neural durations typical of relevant cognitive events.

Supposing that the predicted relationships between stimulus properties and hemodynamic features hold for brief stimuli, high temporal resolution would be needed to reliably detect the subtle timing effects using fMRI. Although the practical temporal effect limit is not known, overall hemodynamic response shifts as small as 28 ms have been reliably detected in visual cortex (Katwal et al., 2013; Rogers et al., 2010; Wang et al., 2017). Timing differences under 100 ms have also been observed in subcortical visual pathway activations (Lewis et al., 2018), thalamic signals during arousal transitions (Setzer et al., 2022), and auditory cortex neural sequences (Li et al., 2024). In most of these studies, characterization of the hemodynamic response was improved with a short TR (repetition time), which allows for more overall samples as well as more fine-grained estimation of the hemodynamic function’s shape (Dowdle et al., 2021; Polimeni & Lewis, 2021). Timing differences on the order of hundreds of milliseconds may be on the edge of standard fMRI paradigms’ resolution, but newer MR sequences and good design choices may allow them to be reliably detected.

Indeed, there has been increasing interest in resolving and using sub-second features of the BOLD signal in recent years, as technology and techniques have both improved (Chen et al., 2023; Li et al., 2024; Marxen et al., 2023; Polimeni & Lewis, 2021; Wittkuhn & Schuck, 2021; Yue et al., 2025).

Another approach that may assist with detecting subtle timing effects is fitting hemodynamic response functions (HRFs) to the collected data. In this approach, a parameterized model is fitted to the data for each participant and condition, effectively filtering the data to reduce noise (Chen et al., 2023; Friston, 2005; Marxen et al., 2023). Parameters such as peak magnitude, peak latency, response latency, and peak dispersion have been shown to constitute meaningful indicators of both within-subject and between-group differences (Lindquist et al., 2009; Wager et al., 2005). Many of these parameters are not orthogonal and can trade off during fitting, so peak latency estimated from the fitted functions offers a more stable and interpretable marker of timing differences (Dux et al., 2006; Tombu et al., 2011; Wager et al., 2005; Yue et al., 2025). HRF fitting approaches can also be combined with other techniques to further increase the signal for temporal metrics.

For example, researchers using ERPs (evoked response potentials) have used resampling techniques, such as bootstrap resampling (Efron & Gong, 1983), to obtain more reliable peak timing estimates, with reduced bias and improved confidence intervals (Bledowski, 2004; Liesefeld, 2018; Miller et al., 1998).

In the present study, we sought to characterize and compare the hemodynamic responses to visual and auditory stimuli that varied in their durations and intensities. In Experiment 1 (n=15), we presented stimuli at three durations (100, 300, and 900 ms) and three intensities (different contrast or volume levels). To best capture the temporal features of the evoked responses, we used a slow event-related design with a rapid TR (625 ms) and then derived key parameters (e.g. peak latency and magnitude) from fitted single-Gamma HRFs, both with and without bootstrap resampling. We hypothesized that peak magnitude would increase with increasing stimulus durations or intensities, whereas only peak latency would increase with increasing stimulus durations. In Experiment 2 (n=14), the stimulus durations were extended (1000, 1200, and 1800 ms) to increase the evoked signal. This change also aligned our design more closely with paradigms that use overlapping evoked signals, which are frequently used in fMRI studies (Dux et al., 2006; Serences, 2004; Yue et al., 2025). We hypothesized the same pattern of results as before, with peak latency uniquely sensitive to stimulus duration changes.

## 2. Experiment 1

In this first experiment, we sought to characterize the effects of parametrically varied visual and auditory stimuli on evoked hemodynamic responses. Stimulus duration and intensity each had three levels, creating a 3×3 design. We hypothesized that, provided noise did not overwhelm our signals, intensity and duration changes would affect observed BOLD magnitude, but only duration changes would affect peak timing (Henson et al., 2002).

Canonical Gamma-shaped hemodynamic response functions (HRFs) were used for formalizing predictions about expected responses and for fitting the empirical data. Parameters derived from these fits were used for standard statistical analyses as well as a bootstrap-based approach inspired by EEG timecourse analyses (Bledowski, 2004; Efron & Gong, 1983; Liesefeld, 2018).

A preview is worth providing here: Experiment 1’s results were suggestive of the hypothesized effects, but they were also inconclusive. The results informed the design of Experiment 2, which provided stronger evidence to address—and support—our primary hypotheses.

### 2.1. Materials and Methods

#### 2.1.1. Experimental procedures

Sixteen members (9 females, 14 right-handed, mean (±SD) age = 23±2.42 years) of the National University of Singapore (NUS) community were recruited for the study, which took place at the Yong Loo Lin School of Medicine’s Clinical Imaging Research Centre (CIRC). Written informed consent was obtained from all participants (for Experiments 1 and 2), in accordance with a protocol approved by the NUS Institutional Review Board. Participants were instructed to avoid caffeine intake in the 24 hours preceding their scanning session. They were paid 40 SGD for their participation.

Participants completed a 70-minute slow event-related main task, conducted across nine 390s runs. Each run began with a 20s fixation period and concluded with a 10s fixation period, with 18 visual trials and 18 auditory trials presented in the interim. Visual trials began approximately every 20^th^ second, with the precise onset jittered from this time by up to 625ms. This jitter reduced participant expectation of event onsets (including based on rhythmic scanner noise) and potentially increased our temporal resolution through subsampling. Auditory trials were presented with similar timing and jitter, save they began 30s into each run. Visual and auditory trials were thus interleaved, with an approximately 10s fixation period between stimuli.

The experiment was presented on a MacBook Air (OS 10.12.1) running PsychoPy v1.83.01 (Pierce, 2007) and connected to a 32-inch LCD monitor (NordicNeuroLab, Bergen, Norway) running at 60 Hz. Each visual trial consisted of a circular contrast-reversing checkerboard (flicker rate: 10Hz) that was 20.7 degrees wide. Participants viewed the stimuli in a supine position via an angled mirror located 12 cm above the eyes. In turn, the distance between this mirror and the LCD monitor was 179 cm. There were 9 possible trial conditions defined by the conjunction of three stimulus durations (100ms, 300ms, and 900ms) and three stimulus intensities (1% contrast, 10% contrast, and 100% contrast). These contrast levels were chosen because increases in visual stimulus intensity by powers of 10 evoke approximately linear increases in peak BOLD signals (Goodyear & Menon, 1998). Each condition combination was repeated twice per run in a random sequence, resulting in each subject experiencing a total of 18 visual trials per condition combination.

Each auditory trial consisted of a repeating sequence of two pure tones (440Hz and 550Hz, with a cycling rate of 10Hz) delivered over MR-compatible pneumatic earphones. Similar to the visual trials, there were a total of 9 possible trial conditions defined by the conjunction of three stimulus durations (100ms, 300ms, and 900ms) and three stimulus intensities (10% volume, 30% volume, and 90% volume). Each trial condition was repeated twice per run in a random sequence, resulting in each subject experiencing a total of 18 auditory trials per condition. To ensure that subjects remained attentive throughout the duration of each scan, subjects responded to each auditory stimulus by pressing a button on an MR-compatible button box (Current Designs, Inc.; Philadelphia, PA, USA) using their right index finger. Eye movements and closures were also monitored using an MR-compatible long-range Eyelink 1000 Plus eyetracker camera (SR Research; Ottawa, Ontario, Canada).

Two functional localiser runs (236s each) followed the main task. Contrast-reversing checkerboards (flicker rate: 10Hz) and oscillating pure tones (440Hz and 550Hz, with a cycling rate of 10Hz) were presented in alternating blocks of 16s separated by 12s fixation periods between blocks.

All but one participant completed the above session plan. This participant completed only six of the planned nine runs, and her data were excluded from further analysis. The final sample therefore included 15 participants.

#### 2.1.2. MRI data acquisition and pre-processing

MRI data were acquired using a Siemens 3T MAGNETOM Prisma MRI scanner (Siemens, Erlangen, Germany) and a 32-channel head coil. Functional scan parameters were adapted from those used for the Human Connectome Project (HCP) and were chosen to ensure that full-brain coverage, including the cerebellum, was achieved for each participant.

A 3D high-resolution (1 x 1 x 1 mm) T1-weighted magnetisation-prepared, rapid acquisition gradient echo (MPRAGE) pulse sequence was used to obtain whole-brain anatomical images for each participant and normalize individual subject data to a standard space. 128 1-mm thick contiguous sagittal slices (0.5 mm skip; 1 x 1 mm in-plane resolution) were acquired with a repetition time (TR) of 2300 ms, an effective echo time (TE) of 2.22 ms, a flip angle (FA) of 8°, and a 260 x 260 mm field of view (FOV).

Functional MRI data were acquired using a multiband echoplanar imaging (MB-EPI; CMRR release R2015) sequence with a MB acceleration factor of 8 (Demetriou et al., 2018). 378 whole-brain images were obtained for each localiser run while 624 images were acquired for each task run. T2*-weighted images were acquired using a TR of 625 ms, a TE of 33.2 ms, and a FA of 50°. 64 interleaved slices (thickness = 2.5 mm, no gap) were collected using a 220 x 220 mm FOV (imaging matrix = 64 x 64). The voxel size was thus 2.5 x 2.5 x 2.5 mm^3^.

Three task runs and one localizer run were discarded due to technical difficulties during data acquisition. The remaining task and localiser data were pre-processed using a previously published pre-processing pipeline (Li et al., 2019; Kong et al., 2019). The initial four images from each run were removed to aid with BOLD signal stabilisation. Motion correction using FSL’s MCFLIRT was then applied such that runs with more than 50% of the frames removed were discarded to ameliorate any contributions of head motion. (This step excluded no runs from the present dataset.) FSL’s bbregister function was used for intrasubject registration of the T1 anatomical images to the T2*-weighted images. The best run for each subject was used as the registration file across all functional runs. Subsequently, motion parameters and their derivatives were regressed out using global signal regression and censored frames were interpolated (Power et al., 2014). Linear trends were removed and the images were spatially smoothed using an isotropic Gaussian kernel of 6 mm (FWHM).

Finally, the images were down sampled and projected to volumetric MNI 2-mm space. These volumetric outputs were subsequently submitted to further analysis in using FSL Release 6.0 (Jenkinson et al., 2012).

#### 2.1.3. Region of interest (ROI) definitions

To account for between-subject differences in BOLD response locations, we sought to define regions-of-interest (ROIs) from each participant’s functional localisers. Each localiser run was first submitted on a per-subject basis to a first-level analysis in FEAT. Stimulation and fixation blocks were modelled using boxcar functions. Model regressors were created by convolving these boxcar functions with a single gamma function (SD = 3s, mean lag = 6s).

The first-level localiser analyses were next combined in a fixed-effects higher-level analysis for each modality and participant, with voxel clustering set at *z >* 3.1 and *p <* .05.

Visual stimulation yielded robust activation clusters in the medial posterior occipital lobes of both the left and right hemispheres. Across participants, there was considerable variation in the spatial location of peak voxel activity. We created per-subject spherical ROI masks with a maximum radius of 6mm. Using FSLeyes, voxels contiguous with the peak were selected within the left and right primary visual cortex. These ROIs were constrained by the anatomical boundaries of primary visual cortex (BA 17 and 18) as defined by the Juelich Histological Atlas (Amunts et al., 2000) with a minimum probability threshold of 1%. The Juelich Histological Atlas is probabilistic; each ROI achieved at least 60% likelihood (left hemisphere, 82% in the right) of overlapping with primary visual cortex.

Functional localization was less successful for auditory localizer runs. Only 6 participants evidenced clusters in the lateral temporal lobes at the chosen threshold of *z >* 3.1 and *p <* .05. Consequently, auditory ROIs were constructed from atlas locations at the group level. ROI masks for the left and right primary auditory cortex were defined from the parcellation of sensory cortices in the Juelich atlas (Eickhoff et al., 2005). The resulting masks demonstrated a substantial degree of overlap with the few clusters of auditory cortex activity observed from the functional localizers. For comparison purposes, such atlas-based ROIs were also defined for primary visual cortex.

#### 2.1.4. Event-related timecourses

Raw BOLD signal values were extracted from each subject’s pre-processed 4-D functional data. These values were averaged across the voxels within each ROI. The resulting timecourse vectors were imported into MATLAB 2018b (Mathworks Inc., Natick MA) for temporal alignment and segmentation. Visual and auditory stimulus presentation times were extracted from scanner logs and converted to the volume of each event onset. Each region-specific BOLD signal vector was segmented into a 20 s event-related timecourse, starting 5 s prior to event onset. Each timecourse terminated 15 s after the event, thereby allowing the hemodynamic response to unfold without contamination from subsequent events or their anticipation.

Raw signal values were next converted into percent signal change values for each event-related time-course. Due to unexpectedly high cross-contamination (e.g. auditory responses in primary visual cortex), the baseline for each event was defined as the measured BOLD signal 1.25s after event onset. This point was chosen because preceding impulses would have fully dissipated, while responses to the present event had not yet emerged. Event-related time-courses were averaged across hemispheres because stimuli had been presented across visual hemifields or binaurally. Responses across hemispheres generally looked similar.

#### 2.1.5. Expected responses from Gamma-based hemodynamic models

For visualization purposes, we generated expected hemodynamic responses for each experimental condition combination. Stimulus durations and intensities were modelled via boxcar functions, with intensity levels (1%, 10% and 100% contrasts; 10%, 30% and 90% volume) coded as linearly-increasing neural activation strengths (i.e., 1, 2, 3). These boxcar functions were then convolved with a canonical hemodynamic response function (Friston et al., 1995) to produce the predicted hemodynamic response curves. As implemented in SPM, the canonical HRF is a double-Gamma function with six parameters (response delay, undershoot delay, response dispersion, undershoot dispersion, ratio of response to undershoot, onset) and a kernel length. For both visualization and function fitting, we used a single-Gamma function by setting the response-to-undershoot ratio to infinity, effectively eliminating the canonical HRF’s undershoot component. The starting values of the other parameters followed the SPM defaults. Predictions from the forward models are presented alongside the relevant empirical results.

#### 2.1.6. Fitting Gamma-based hemodynamic models to empirical data

Parameterised canonical hemodynamic response functions were also used for visualization and statistical analysis of the empirical data. Event-related average BOLD time-courses were first aggregated by condition combination. For visualization, each aggregated sample included all participants, which amounted to approximately 4800 discrete events (15 subjects x 9 runs x 2 ROIs x 9 conditions x 2 events per condition). For standard statistical analyses, aggregates were created for each participant separately. Bootstrap-based statistical analysis used aggregates created by sampling each participant’s data with replacement; the motivation and procedures for this approach are detailed below.

For each aggregated data set, the best-fit canonical hemodynamic response function (adapted from the double Gamma SPM implementation (Friston et al., 1995)) was found using a non-linear least squares algorithm implemented in MATLAB 2018b (Mathworks Inc., Natick MA). As the ratio of response to undershoot was set to infinite, the response function was a single Gamma with four free parameters: peak amplitude, response onset, peak delay, and peak dispersion. This approach was similar to Wager et al. (2005)’s, save the addition of peak dispersion as a free parameter. Its inclusion produced modelled hemodynamic response curves that were better aligned with the event-related average time-courses, particularly in the response’s falling phase.

The obtained parameter estimates were used to generate hemodynamic response curves. Many of the HRF parameters are not orthogonal and can trade off during fitting (Lindquist et al., 2009; Wager et al., 2005), so we instead derived key parameters from the generated curves themselves, thereby obtaining more stable and interpretable markers for statistical analyses (Dux et al., 2006; Tombu et al., 2011; Wager et al., 2005; Yue et al., 2025). Peak magnitude and peak latency were defined as the maximum BOLD percent signal change and its corresponding timepoint. Response onset latency was defined as the time at which the hemodynamic response reached 10% of the peak amplitude. Peak dispersion was defined as the full-width half-max (FWHM) value of the curve. Group means, as well as within-subject standard deviations and 95% confidence intervals (CIs), were subsequently calculated for all four measures (Cousineau, 2005).

#### 2.1.7. Bootstrap resampling and statistical analyses

Conventional inferential statistics rely on metric estimates from individual subject data, with each subject’s measures assumed to be equally reliable (Suchow et al., 2013). Neuroimaging measures, however, exhibit considerable variance across timepoints and participants, rendering it challenging to derive good estimates of hemodynamic response onset and peak latency—even after using the Gamma-based fitting procedures described above. A potential solution comes from EEG analyses, in which evoked response potential timing is also important but difficult to estimate (Liesefeld, 2018). Group-averaged timecourses increase the signal-to-noise ratio, thereby allowing for the estimation of reliable sample parameters (Asplund et al., 2010). Variance of the estimates is obtained through resampling.

Here we used bootstrap resampling to estimate parameter means and variances, in addition to calculating traditional by-participant estimates. For our bootstrap procedure, 10,000 “pseudo-samples” of equivalent size to the original dataset were constructed by sampling by-participant data (mean event-related timecourses for each condition combination) at random and with replacement (Efron & Gong, 1983). That is, a given bootstrap sample had data for each condition combination from the same set of participants, allowing us to calculate within-subject estimates for differences between parameters (e.g. peak latency for 100% contrast, 100 ms duration versus 100% contrast, 900 ms duration). For each bootstrap sample, the best-fit canonical hemodynamic response function was found using a non-linear least squares algorithm, which was then used to generate hemodynamic response curves from which the bootstrapped parameter values were obtained.

Statistical analysis of bootstrapped and non-bootstrapped parameter estimates followed the same procedures to enable direct comparisons. For each parameter, we computed means, standard deviations (SD), standard errors (SE), and 95% confidence intervals (CIs) across all nine study condition combinations for each stimulus modality. We also report mean differences and their CIs, using these intervals as our primary inferential tool, rather than performing traditional null-hypothesis significance tests (NHST). This estimation-based approach emphasizes effect size and precision, an important focus when small but meaningful timing differences are expected. Our approach aligns with recent recommendations to prioritize effect estimation over binary significance testing, particularly in contexts where precision and interpretability are critical (Claridge-Chang & Assam, 2016).

Unless stated otherwise, figures and reported findings in Experiment 1 reflect bootstrapped parameter estimates. Standard parameter estimates were used for calculating effect sizes (e.g., Cohen’s *d*). Figures and reported findings for Experiment 2 reflect conventional by-participant estimates.

### 2.2 Results

#### 2.2.1. Visual stimulus duration and intensity effects were consistent with predicted HRFs

The simulated BOLD responses illustrated the expected effects of stimulus intensity and duration. When stimulus duration increased but intensity was held constant, both peak magnitude and peak latency increased (Figure 1, top row). The empirical results were broadly consistent with these simulated effects. At maximal stimulus contrast (100%), increases in visual stimulus duration led to increases in both peak magnitude and peak latency of the fitted hemodynamic responses (Figure 1, bottom row). Each increase in stimulus duration led to a statistically significant increase in average peak magnitude, which doubled across the tested stimulus duration range (large effect size, d = 1.44). Pairwise changes in peak latency, however, were too variable to yield statistically significant effects. The substantial and significant 0.72 s increase (95% CI [0.30 1.18], d = 0.74) in peak latency across the 100 ms to 900 ms stimuli accorded with predictions, but its variability was likewise large. Note that in the confidence interval framework, an effect is considered statistically significant when the 95% CIs do not include zero (Claridge-Chang & Assam, 2016).

**Figure 1.**
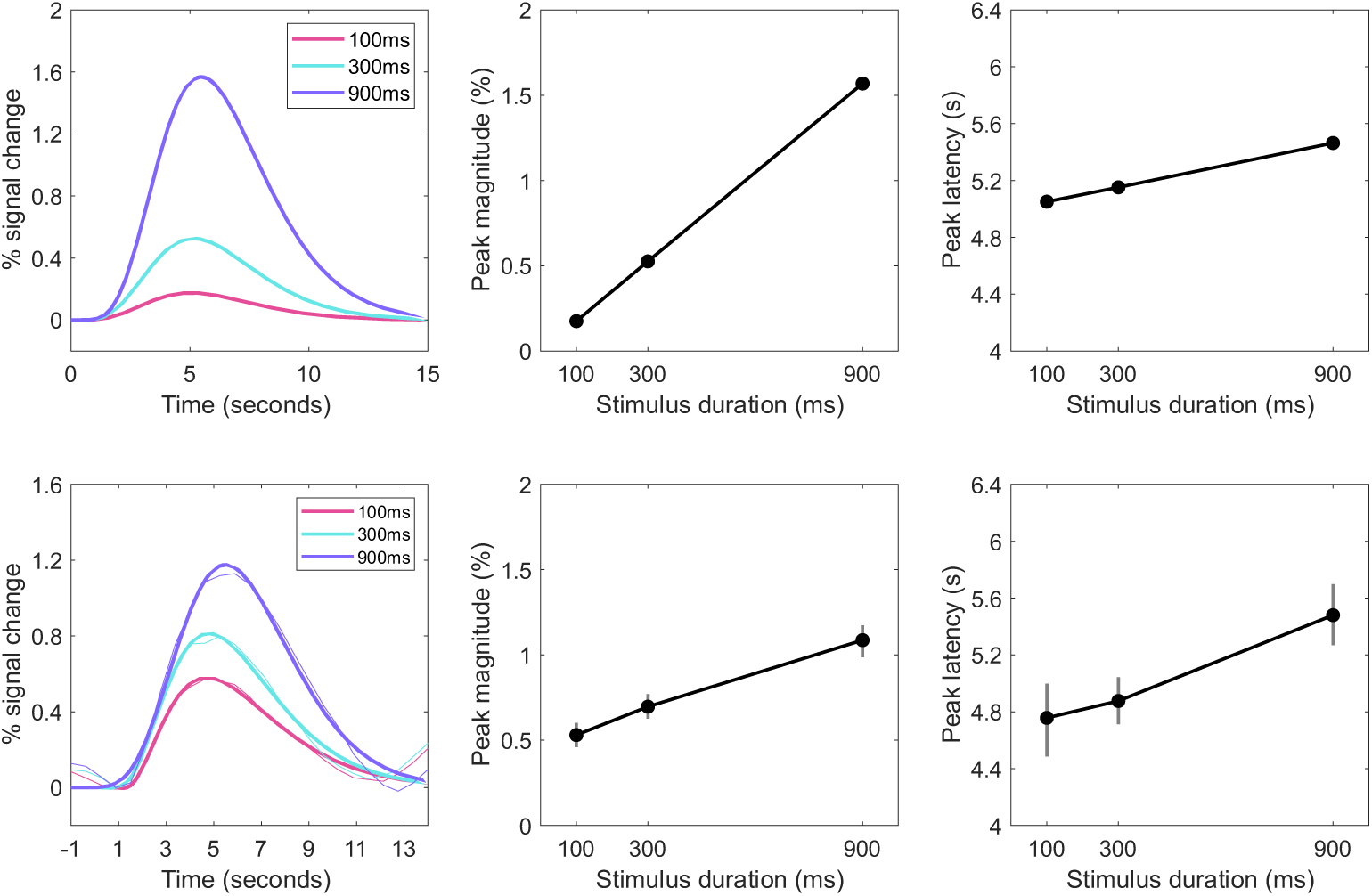
Predicted (top row) and observed (bottom row) effects of stimulus duration on the overall hemodynamic response (left column), peak magnitude (middle column), and peak latency (right column) evoked by maximum-contrast (100%) stimuli in primary visual cortex. Observed hemodynamic responses were obtained by averaging across participants. Observed peak magnitude and peak latency estimates and 95% CIs were derived from bootstrap samples.

Increases in stimulus duration led to increases in both the peak magnitude and peak latency of the event-related hemodynamic response, albeit with high variance in the latter metric.

The simulated BOLD responses also illustrated the expected contrast between manipulations of stimulus duration and intensity. When stimulus intensity increased but duration was held constant, only the magnitude of the hemodynamic response increased (Figure 2, top row). The empirical effects were again broadly consistent with the simulations. At the 900 ms stimulus duration, peak magnitude increased as stimulus contrast increased, doubling across the tested range (large effect size, d = 1.29; Figure 2, bottom row). Peak latency showed only small (0.20 s, 95% CI [-0.59 0.71], d = 0.37) and non-significant changes across the tested range.

**Figure 2.**
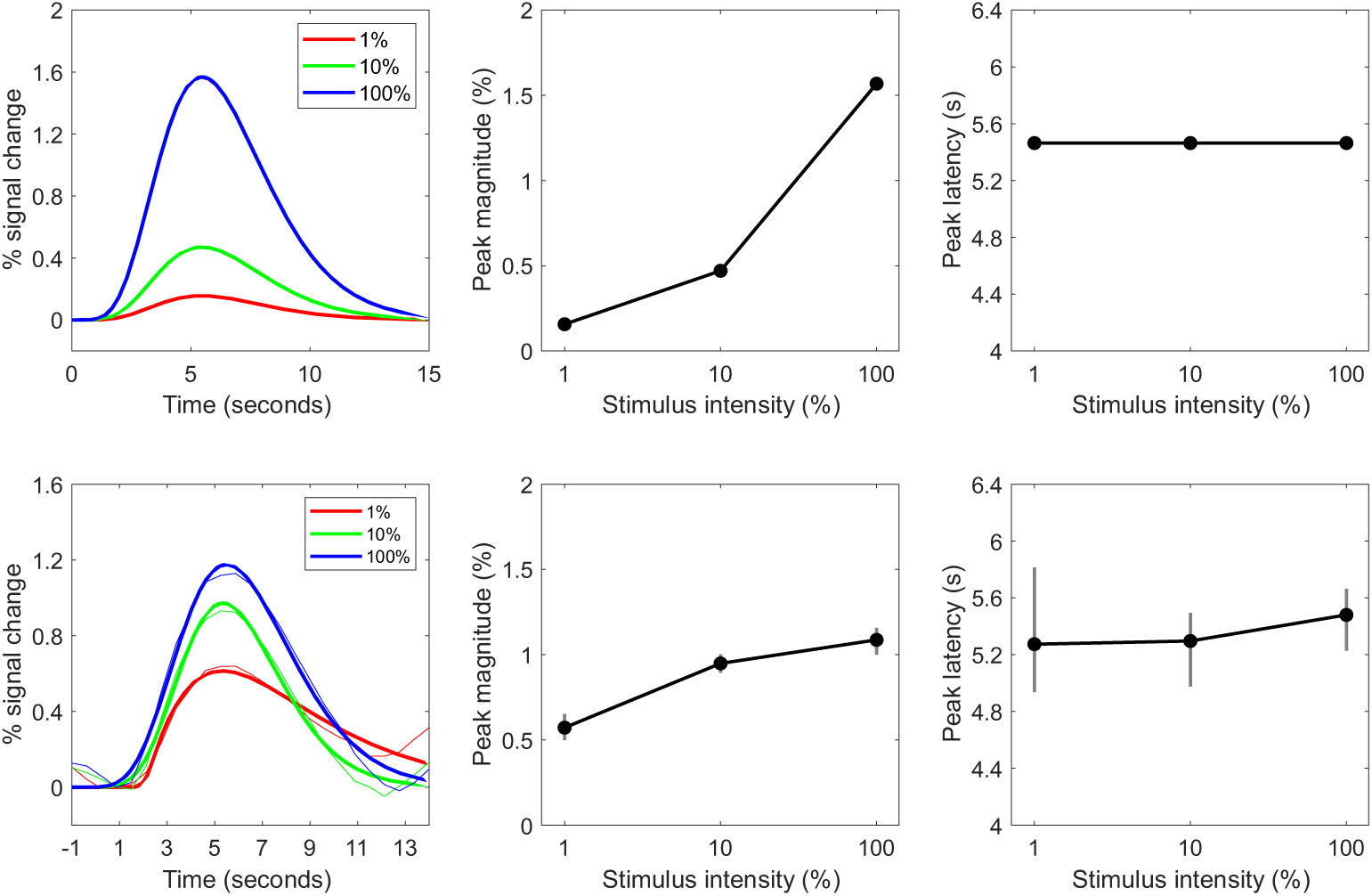
Predicted (top row) and observed (bottom row) effects of stimulus intensity on the overall hemodynamic response (left column), peak magnitude (middle column), and peak latency (right column) evoked by 900 ms stimuli in primary visual cortex. Increasing stimulus intensity (i.e., contrast) led to increases in peak magnitude, but not peak latency, of the event-related hemodynamic response.

Notably, the standard analytical approach produced a mean peak latency of 6.07 s (SD = 1.66, 95% CI [5.23, 6.92]) for the 1% contrast condition, as compared to 5.47 s (SD = 0.83, 95% CI [5.05, 5.90]) for the 10% contrast condition. Inspection of the individual fits revealed peak latencies in excess of 6 s for 5 of 15 participants in the 1% contrast condition, for which overall activation was low. This situation illustrates the difficulty of deriving robust temporal parameters in the presence of noise that overwhelms the signal (Liesefeld, 2018). The bootstrap approach is a potential remedy to this problem. Indeed, the bootstrapped peak latency estimates for 900 ms stimuli differed by just 0.2 s across the three levels of stimulus contrast, perhaps reflecting greater metric stability. Regardless, metrics derived from either approach were not consistent with increasing peak latency with increasing stimulus contrast. Parameter estimates and their variances for each condition combination can be found in the Supplementary Materials as csv tables.

#### 2.2.2. Latency estimates were highly variable at lower visual stimulus intensities

Peak magnitude was sensitive to manipulations of stimulus duration at lower contrast levels (Figure 3, bottom row; Figure 4). Similarly, peak magnitude reflected contrast manipulations with shorter-duration stimuli (Figure 3, top row; Figure 4). For example, the difference between 1% and 100% contrast for 300 ms stimuli had a large effect size (d = 1.30), comparable to the results for 900 ms stimuli.

**Figure 3.**
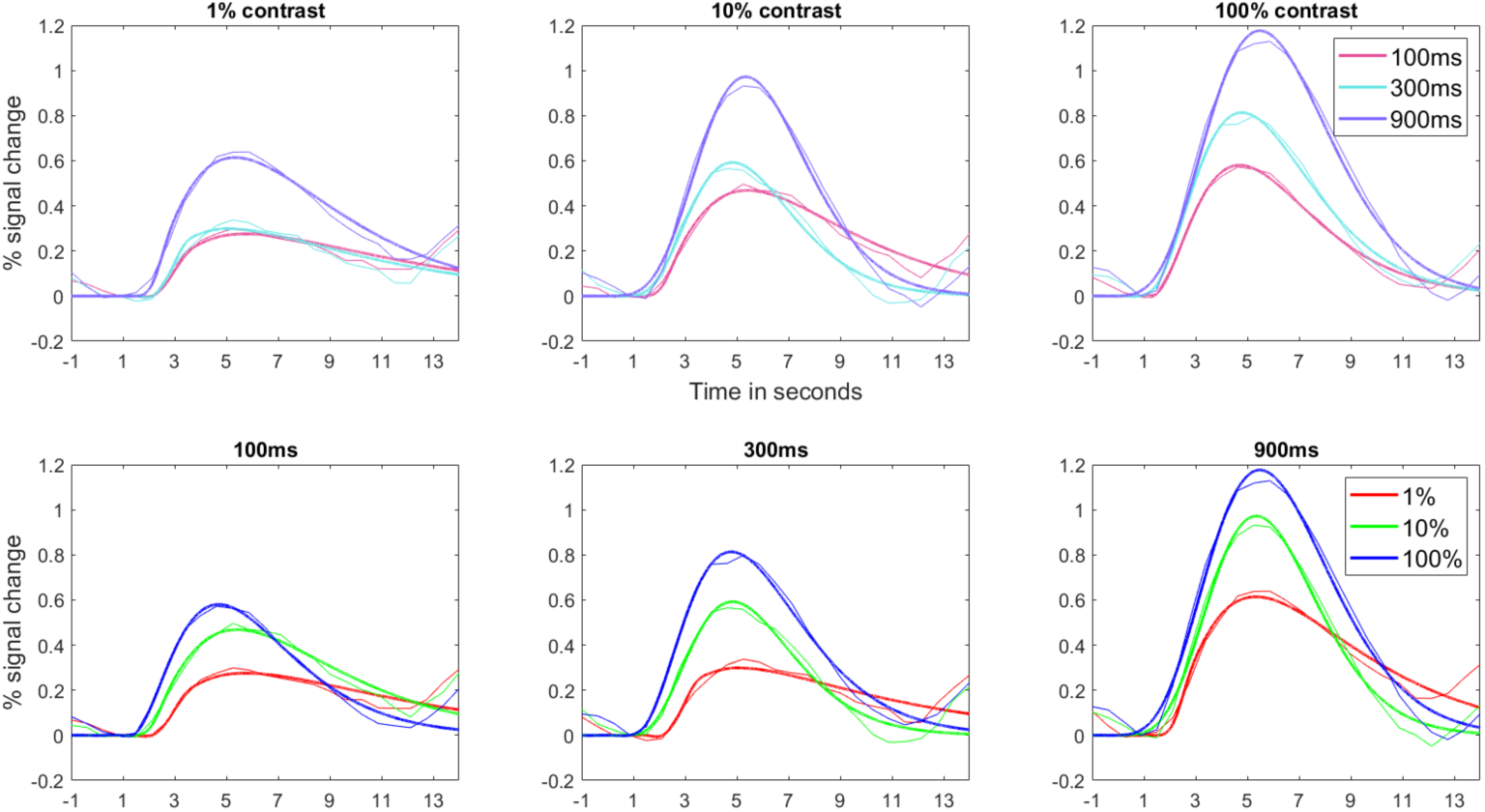
Event-related hemodynamic responses in primary visual cortex across increasing levels of stimulus intensity at each level of stimulus duration (top row) and increasing levels of stimulus duration at each level of stimulus intensity (bottom row).

**Figure 4.**
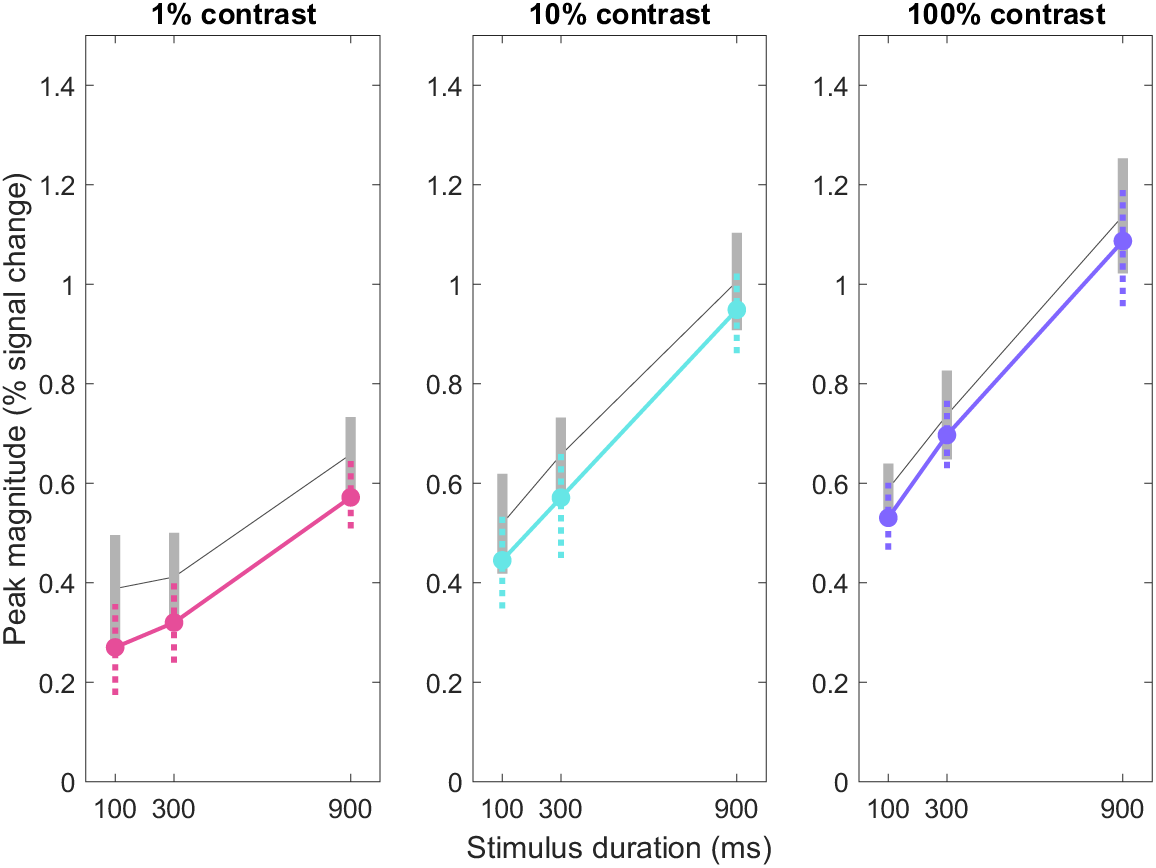
Peak magnitude estimates across visual stimulus durations and contrasts. Increases in stimulus durations were associated with increases in peak magnitudes of the corresponding event-related hemodynamic responses at each level of visual stimulus contrast. Bootstrapped estimates and error bars are displayed in colour; non-bootstrapped estimates and error bars are plotted in grey. Error bars represent 95% CIs. Significant increases in mean peak magnitude can be observed between responses to 300ms and 900ms stimuli at all levels of stimulus contrast.

At lower levels of stimulus contrast, peak latency differences—or the lack of them— could not be established (Figure 3, Figure 5). In particular, the observed responses in the 1% contrast condition often had very low amplitudes or deviated from expected HRF shapes.

**Figure 5.**
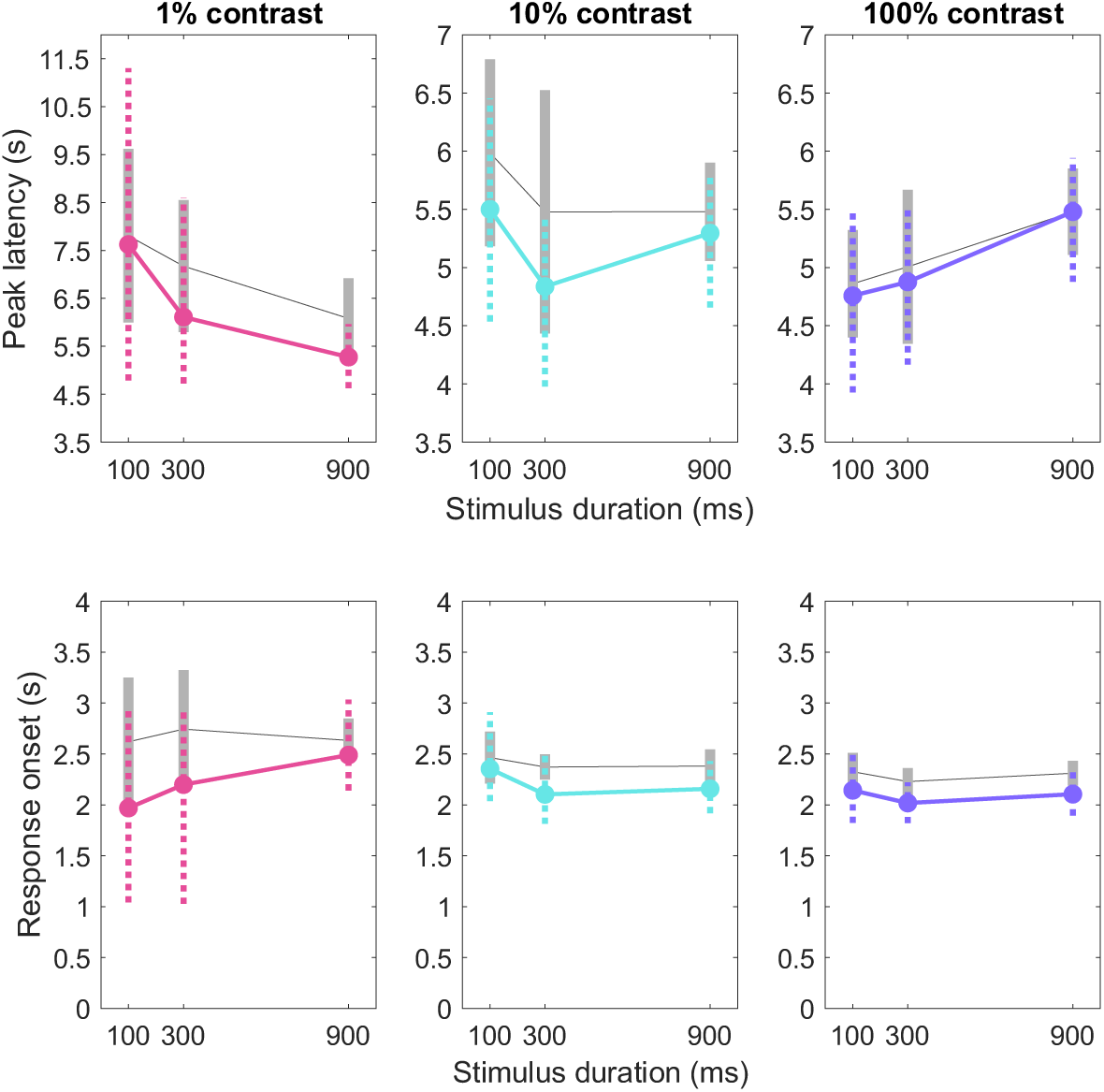
Peak latency and response onset estimates across visual stimulus durations and contrasts. Bootstrapped estimates and error bars are displayed in colour; non-bootstrapped estimates and error bars are plotted in grey. Error bars represent 95% confidence intervals of the mean. There were negligible effects of stimulus duration on the response onsets (bottom row) of the corresponding event-related hemodynamic responses at each level of stimulus contrast.

Accordingly, the variance for these estimates was high. Bootstrap resampling did not appreciably reduce this problem in such low-signal conditions.

#### 2.2.3. Auditory stimulus duration and intensity effects were consistent with predicted HRFs

The empirical results were broadly consistent with the forward model predictions (Figure 6, top row), albeit less convincingly so than in the visual domain. When stimulus volume was held constant at 90% of the maximum, increases in auditory stimulus duration led to increases in both peak magnitude and peak latency of the fitted hemodynamic responses (Figure 6, bottom row). Across the tested range (100 to 900 ms), peak magnitude increased by 0.23% (SE = 0.06, 95% CI [0.10 0.35], d = 1.03) and peak latency increased by 0.27 s (SE = 0.08, 95% CI [0.10 0.42], d = 0.55).

**Figure 6.**
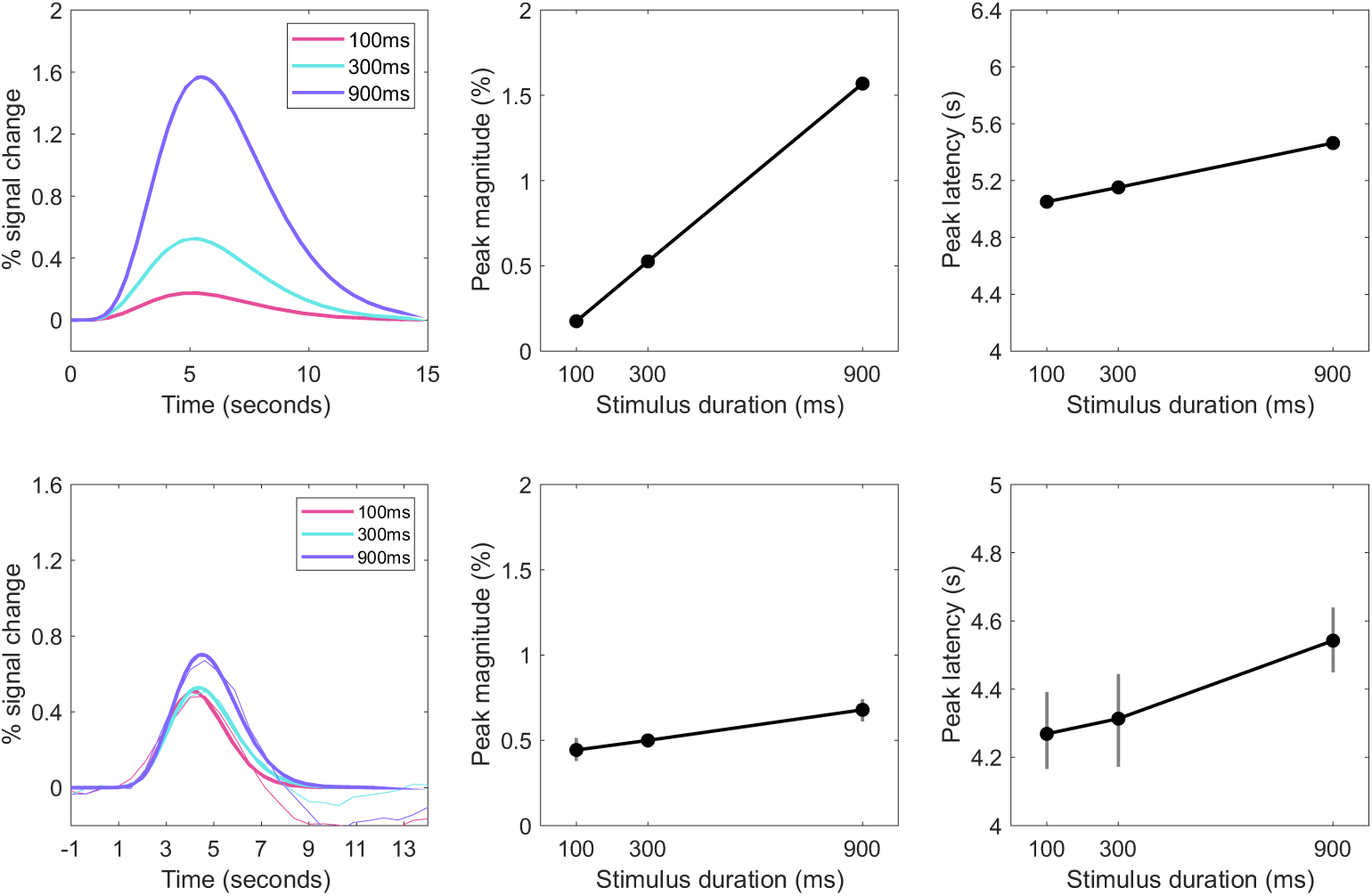
Predicted (top row) and observed (bottom row) effects of stimulus duration on the overall hemodynamic response (left column), peak magnitude (middle column) and peak latency (right column) evoked by stimuli at the maximum tested volume (90%) in primary auditory cortex. Observed peak magnitude and peak latency estimates and 95% CIs were derived from bootstrap samples.

Conversely, when auditory stimulus intensity increased but duration was held constant at 900 ms, peak magnitude moderately increased while peak latency showed no evidence of an increase. Indeed, across the tested range, peak magnitude increased by 0.23% (SE = 0.05, 95% CI [0.11 0.34], d = 0.99), but peak latency (non-significantly) decreased by 0.24s (SE = 0.08, 95% CI [−0.38 −0.08], d = −0.17). Parameter estimates and their variances for each condition combination can be found in the Supplementary Materials as csv tables.

#### 2.2.4. Latency changes were small at lower auditory stimulus intensities

At lower stimulus intensities, peak magnitude was sensitive to manipulations of auditory stimulus duration (Figure 8, bottom row; Figure 9). Similarly, peak magnitude reflected intensity (volume) manipulations even with short-duration stimuli (Figure 8, top row; Figure 9). Some peak magnitude differences were less pronounced than their visual counterparts. That difference could reflect the stimulus intensity factors used (10x for visual, 3x for auditory) or the influence of background acoustic noise on baseline auditory cortex activation as well as perception.

**Figure 7.**
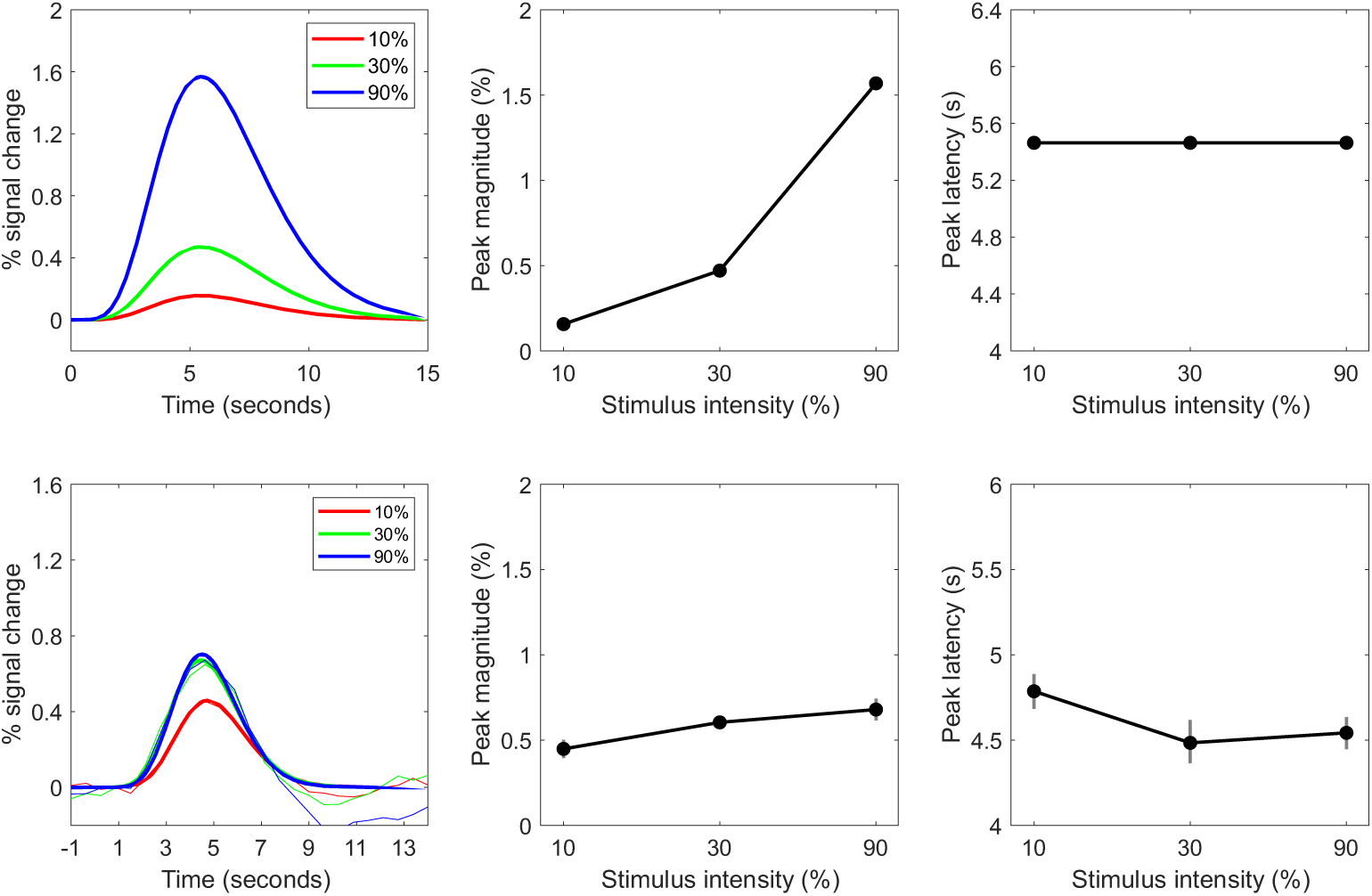
Predicted (top row) and observed (bottom row) effects of stimulus intensity on the overall hemodynamic response (left column), peak magnitude (middle column) and peak latency (right column) evoked by 900ms stimuli at primary auditory cortex. Observed peak magnitude and peak latency estimates and 95% CIs were derived from bootstrap samples. Increasing stimulus intensity (i.e., volume) leads to increases in the peak magnitude, but not the peak latency, of the event-related hemodynamic response.

**Figure 8.**
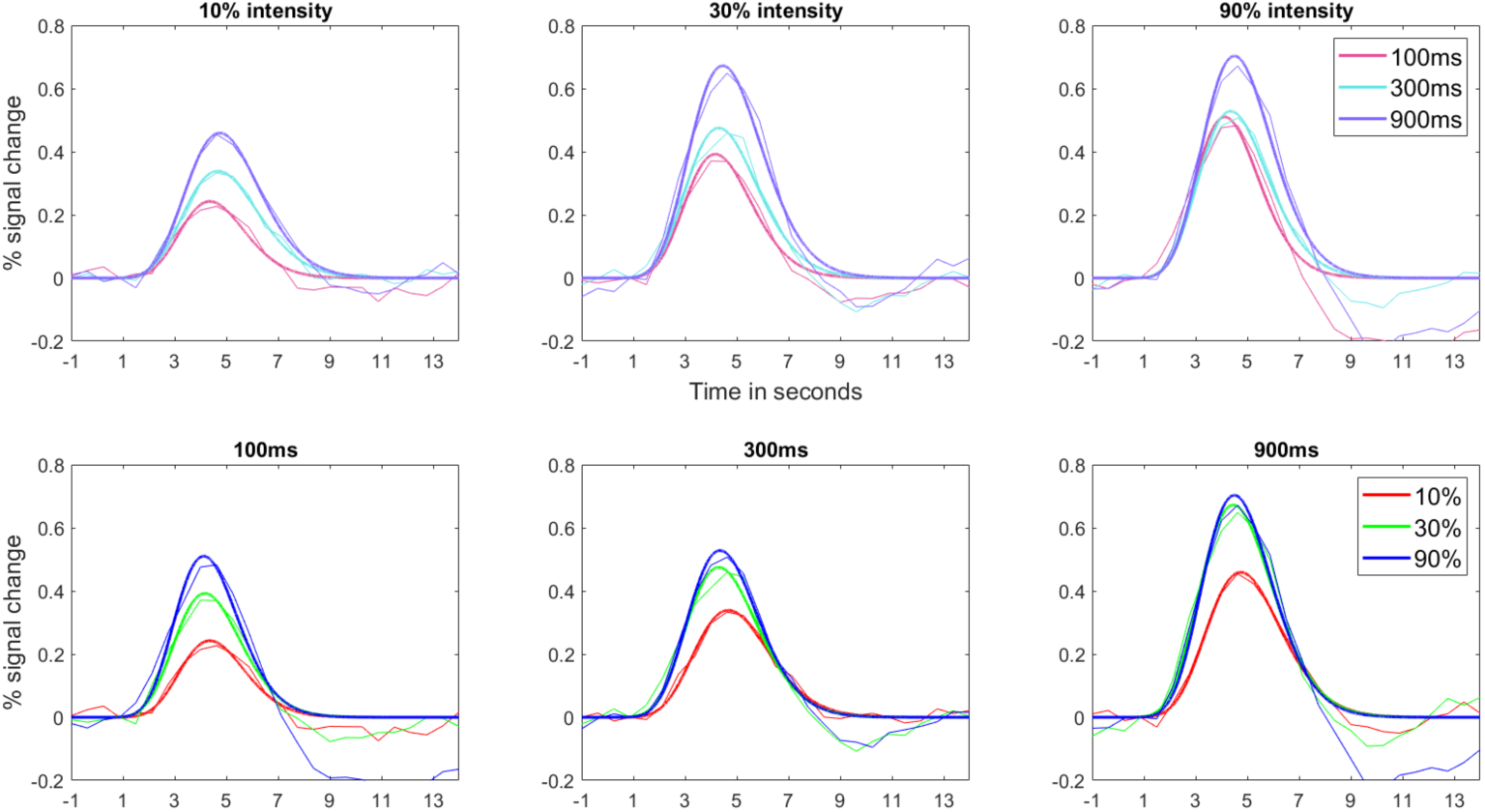
Event-related hemodynamic responses in primary auditory cortex across increasing levels of stimulus intensity at each level of stimulus duration (top row) and increasing levels of stimulus duration at each level of stimulus intensity (bottom row).

**Figure 9.**
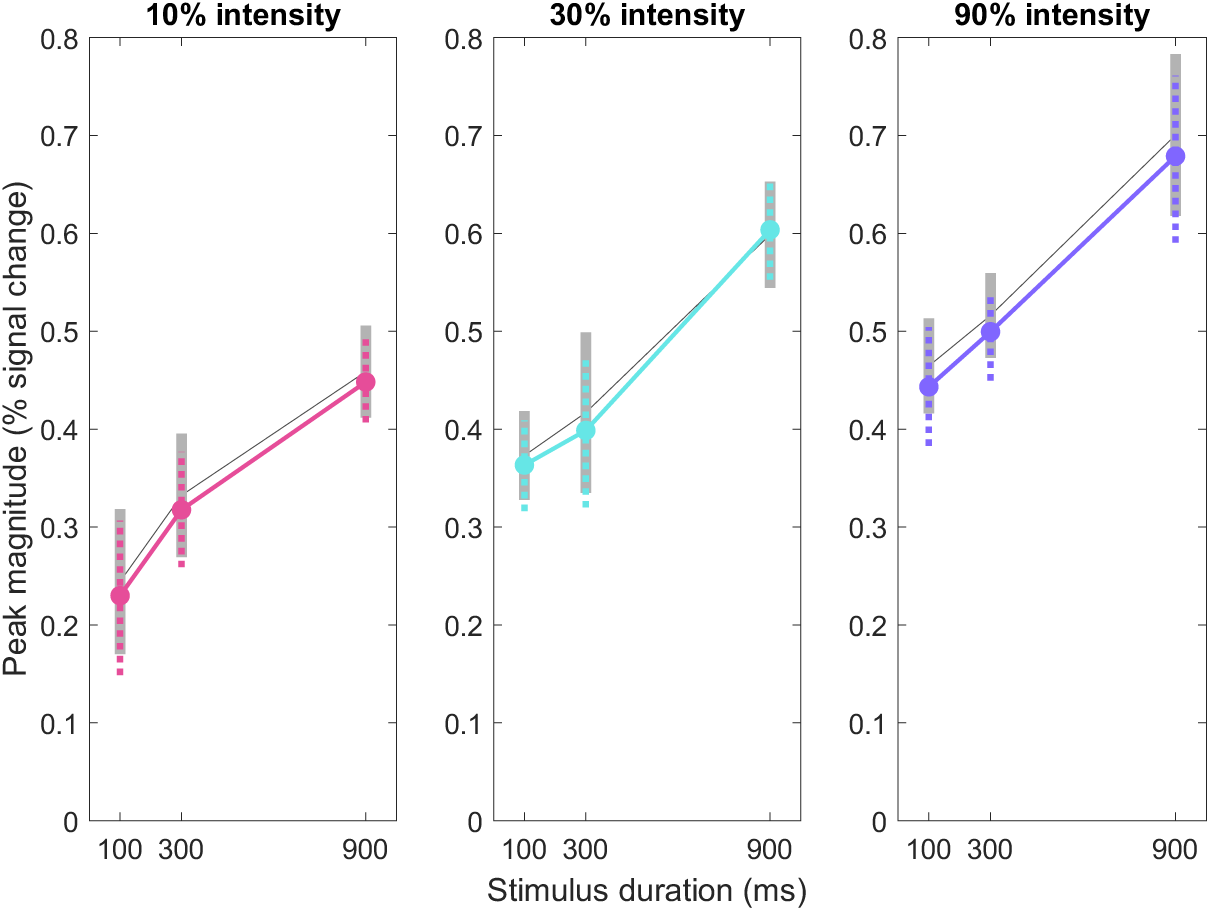
Peak magnitude estimates across auditory stimulus durations and contrasts. Increases in stimulus duration were associated with suggestive but non-significant increases in peak magnitudes of the corresponding event-related hemodynamic responses at each level of auditory stimulus contrast. Bootstrapped estimates and error bars are displayed in colour; non-bootstrapped estimates and error bars are plotted in grey. Error bars represent 95% confidence intervals of the mean.

Unlike in the visual domain, however, peak latency estimates remained relatively robust even at low stimulus intensity levels (Figure 10). Few individuals showed very low activation, even in the low volume conditions. The bootstrapping procedure appeared to be more helpful here as well, perhaps because its use ameliorated random noise contributions instead of apparent deviations from standard HRFs.

**Figure 10.**
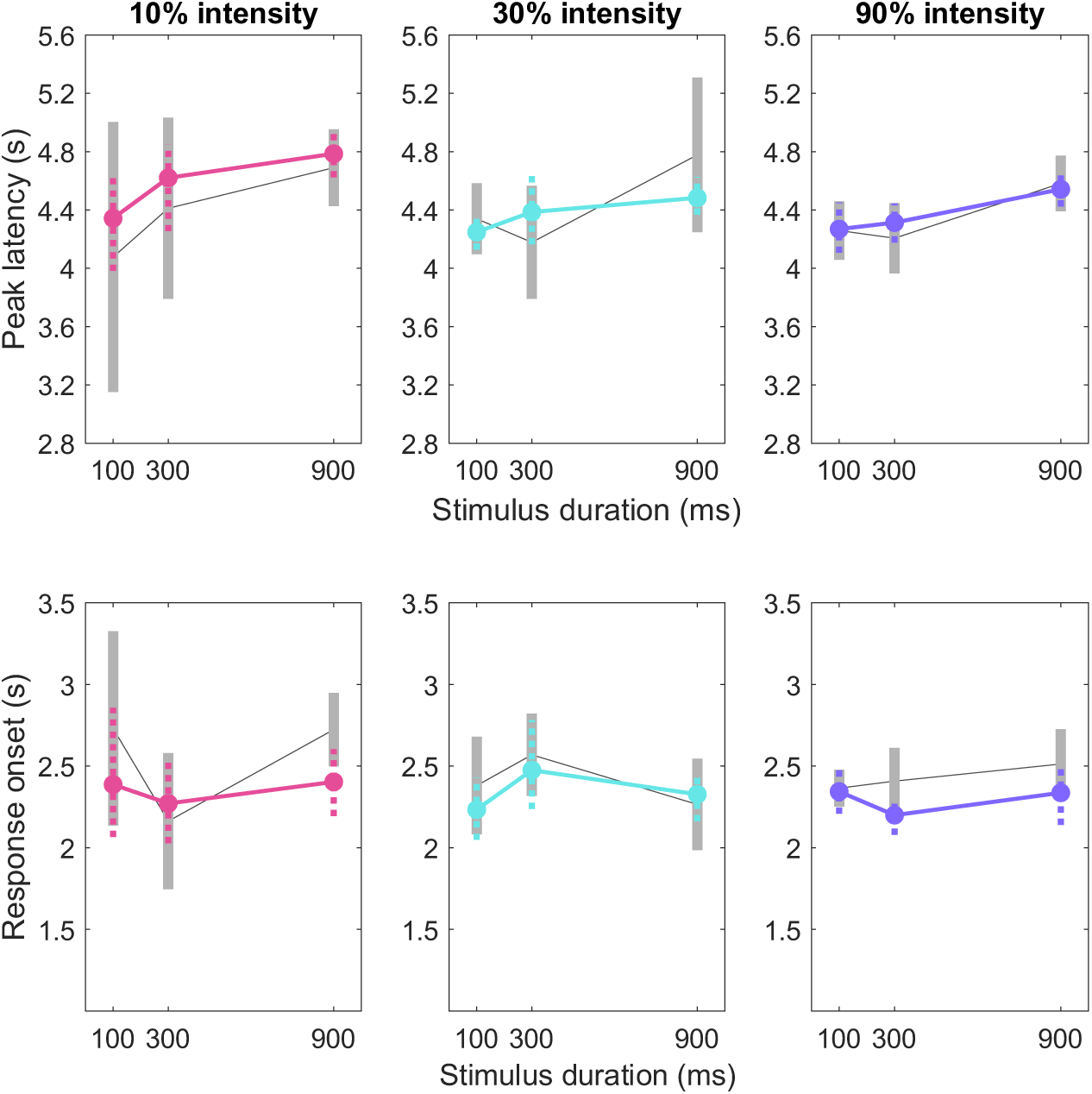
Peak latency and response onset estimates across auditory stimulus durations and contrasts. Increases in stimulus duration led to non-significant but positive trends in mean peak latencies of event-related hemodynamic responses at primary auditory cortex across all levels of stimulus intensity. Bootstrapped estimates and error bars are displayed in colour; non-bootstrapped estimates and error bars are plotted in grey. Error bars represent 95% confidence intervals of the mean. There were negligible effects of stimulus duration on response onset (bottom row) at each level of stimulus intensity.

### 2.3. Experiment 1 Discussion

Experiment 1 investigated whether the BOLD signal’s peak magnitude and peak latency were sensitive to manipulations of stimulus duration and intensity. These parameters were estimated from double-Gamma hemodynamic functions fitted to individual subject data or bootstrap samples. For both auditory and visual stimuli, the results were largely consistent with predictions and previous empirical findings. Foremost, increases in stimulus duration or intensity produced non-linear increases in peak magnitude (Liu & Gao, 2000; Vazquez & Noll, 1998; Wager et al., 2005; Yeşilyurt et al., 2008). In contrast, only increases in stimulus durations appeared to affect peak latency.

For most comparisons of interest, however, substantial variability in parameter estimates (wide 95% confidence intervals) precluded drawing strong inferences about the effects of experimental manipulations on those parameters. In particular, peak latency differences could not be confidently established for stimulus duration changes. Moreover, there remained significant variability in the estimated hemodynamic response parameters at lower stimulus intensities, even when using bootstrap resampling approaches. This result may reflect a limit in the extent to which the canonical HRF can characterise the BOLD responses evoked by the brief periods of visual and auditory stimulation. When analysing such responses, canonical HRF models may demonstrate reduced sensitivity to activation, as rapid signal fluctuations and temporal heterogeneity across voxels lead to responses that are both smaller in amplitude and more variable across trials (Polimeni & Lewis, 2021). The findings informed the design changes in Experiment 2, which prioritized stronger evoked responses and more interpretable parameter estimates.

## 3. Experiment 2

In Experiment 1, brief visual and auditory stimuli often evoked hemodynamic responses with low signal-to-noise ratios, resulting in highly variable parameter estimates. To address this limitation, Experiment 2 included several modifications to improve raw signal quality, allowing us to obtain more reliable parameter estimates without relying on resampling procedures. Foremost, stimulus durations were increased to evoke robust hemodynamic responses, while preserving the small relative differences between stimuli (200 or 600 ms). To improve data quality and boost statistical power, the number of trials per condition was increased and the number of conditions was reduced. A longer TR (1000 ms) was adopted to increase the signal-to-noise ratio of the collected data. Aside from these changes, Experiment 2 retained many of the features of Experiment 1, using the same types of visual and auditory stimuli, preserving the same differences between each level of stimulus duration, and presenting these stimuli at similar intervals within a similar run length. We predicted that duration and intensity changes would both affect peak magnitudes, whereas only duration changes would affect peak latency.

### 3.1. Materials and Methods

#### 3.1.1. Experimental procedures

The procedures were identical to those in Experiment 1 except as noted. Fourteen members from the NUS community (5 female, 13 right-handed, mean (±SD) age = 27.5±2.36 years) were recruited for the study. They were paid 30 SGD for their participation.

Participants underwent a 60-minute slow event-related main task, conducted across 8 runs of 350 s each. Each run consisted of 16 visual trials and 16 auditory trials. The same contrast-reversing and alternating-tone stimuli from Experiment 1 were used.

There were four visual trial conditions. For three of them, the visual stimuli were presented at 100% contrast for 1000 ms, 1200 ms, or 1800 ms. In the fourth condition, the visual stimuli were presented at 10% contrast for 1200ms. Each trial condition was repeated four times per run in a random sequence, resulting in each subject experiencing a total of 32 visual trials per condition over the course of the entire task.

As with the visual trials, there were four auditory trial conditions. For three of them, the auditory stimuli were presented at 90% volume for 1000 ms, 1200 ms, or 1800 ms. In the fourth condition, auditory stimuli were presented at 30% volume for 1200 ms. Each trial condition was repeated four times per run in a random sequence, resulting in each subject experiencing a total of 32 auditory trials per condition over the course of the entire task. To ensure that subjects remained attentive throughout the duration of each scan, subjects responded to each auditory stimulus by pressing a button on an MR-compatible button box (Current Designs, Inc.; Philadelphia, PA, USA) using their right index finger.

Two functional localiser runs (236 s each) followed the main task. Contrast-reversing checkerboards (flicker rate: 10 Hz) and oscillating pure tones (440 Hz and 550 Hz, with a cycling rate of 10 Hz) were presented in alternating blocks of 16 s separated by 12 s fixation periods between blocks.

All but two participants completed the above session plan. As these participants did not complete at least 80% of the planned runs, their data was excluded from further analysis. The final sample therefore consisted of 12 participants.

#### 3.1.2. MRI data acquisition, pre-processing, and ROI definition

The MRI data acquisition approach was the same as in Experiment 1 with the following exceptions. Functional MRI data were acquired using a multiband echoplanar imaging (MB-EPI; CMRR release R2015) sequence with a MB acceleration factor of 6 (reduced from 8 in Experiment 1). 236 whole-brain images were obtained for each localiser run while 350 images were acquired for each task run. T2*-weighted images were acquired using a TR of 1000 ms, a TE of 31.8 ms, and a FA of 50°. 60 interleaved slices (thickness = 2.5 mm, no gap) were collected using a 220 x 220 mm FOV (imaging matrix = 64 x 64). The voxel size was thus 2.5 x 2.5 x 2.5 mm^3^.

All task and localiser data were pre-processed using the same procedures as in Experiment 1. None of the runs were excluded from the dataset during pre-processing. Visual stimulation from the localisers yielded robust activation clusters in the medial posterior occipital lobes of both the left and right hemispheres. Spherical ROI masks were created for each subject with a maximum radius of 6mm. Using FSLeyes, voxels contiguous with the peak were selected within the left and right primary visual cortex. These ROIs were constrained by the anatomical boundaries of primary visual cortex (BA 17 and 18) as defined by the Juelich Histological Atlas (Amunts, Malikovic, Mohlberg, Schormann, & Zilles, 2000).

Auditory stimulation from the localisers also yielded robust activation clusters at the lateral temporal lobes of both the left and right hemispheres for 10 of 12 participants.

Spherical ROI masks (maximum radius of 6 mm) were created for each subject who had robust auditory activation. For the remaining two participants, auditory ROIs were constructed from atlas locations.

#### 3.1.3. Event-related timecourse extraction, Gamma-based hemodynamic models

In Experiment 2, the baseline for each event was defined as the BOLD signal level occurring 1s before event onset. For visualization purposes, event-related time-courses were averaged by modality, collapsing across hemispheres because stimuli had been presented across visual hemifields or binaurally. As in Experiment 1, responses across hemispheres looked similar. Gamma-based hemodynamic response fitting, parameter extraction, and analysis procedures were identical to those from Experiment 1 unless otherwise noted. In particular, because the quality of the timecourses was higher in Experiment 2, we analyzed condition-wise HRF parameters using standard participant-level estimates, without relying on bootstrap resampling.

### 3.2. Results

#### 3.2.1. Peak latency differentiates visual stimulus duration and intensity manipulations

As predicted, increases in visual stimulus durations led to significant increases in both peak magnitudes and peak latencies of the hemodynamic response (Figure 11, middle left; Figure 12). Peak magnitude increased by 0.12 (SD = 0.18, 95% CI [0.01 0.22]) as stimulus duration increased by 200 ms (from 1000 to 1200 ms), and it increased by 0.38 (SD = 0.20, 95% CI [0.26 0.49]) as stimulus duration increased by 600 ms (from 1200 to 1800 ms). The overall peak magnitude increase of 0.50 (SD = 0.16, 95% CI [0.40 0.59]) was similar to the increase observed in Experiment 1 for the same stimulus duration change.

**Figure 11.**
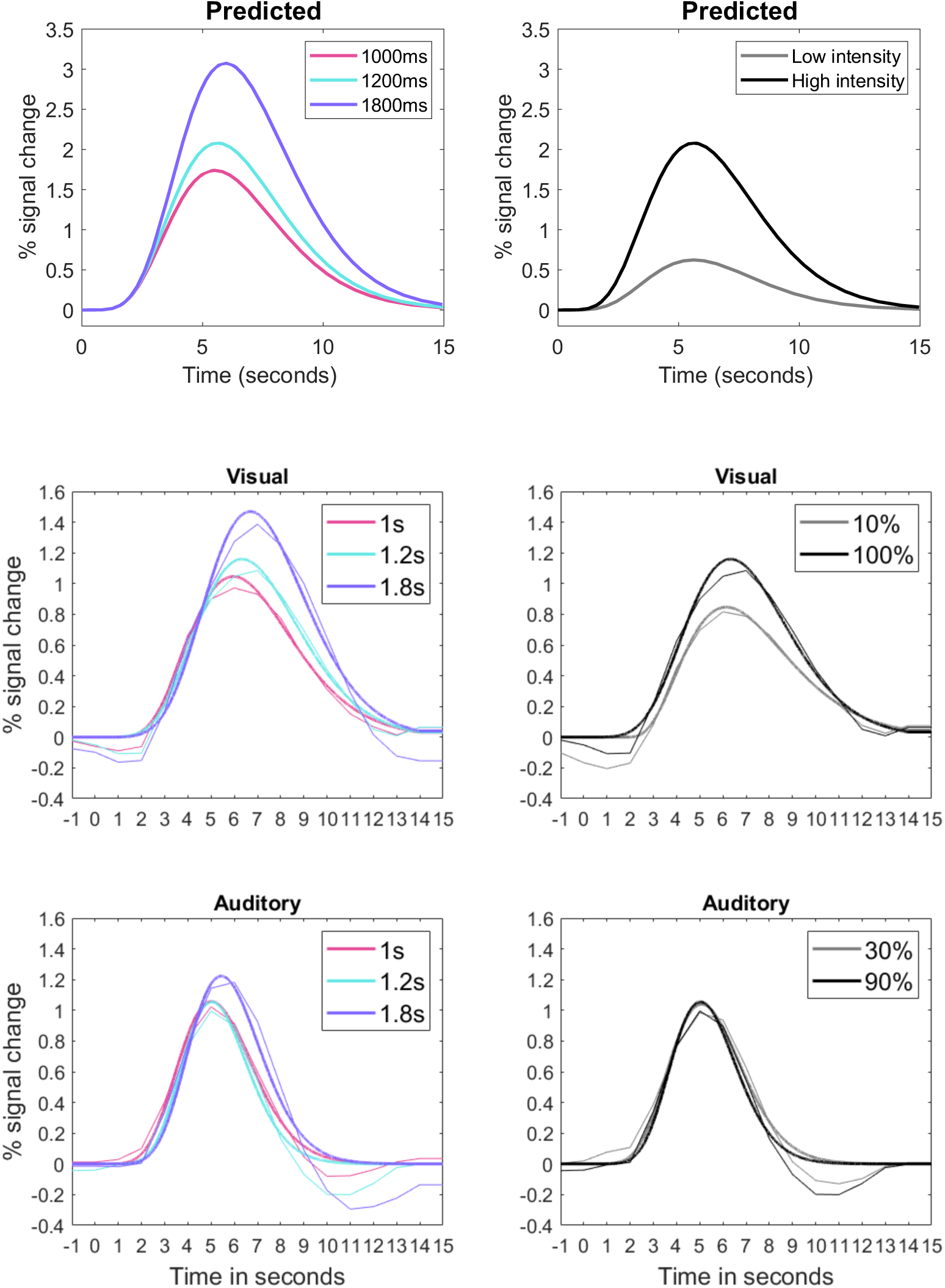
Predicted (top row) and observed event-related hemodynamic responses in primary visual cortex (middle row) and primary auditory cortex (bottom row). Effects of visual and auditory stimulus durations of 1000 ms, 1200 ms and 1800 ms at maximal stimulus intensity (left column) and across two levels of visual and auditory stimulus intensity at the 1200 ms stimulus duration (right column). Note the peak latency changes for duration manipulations—which were clear even with the longer TR—but not for different intensities.

**Figure 12.**
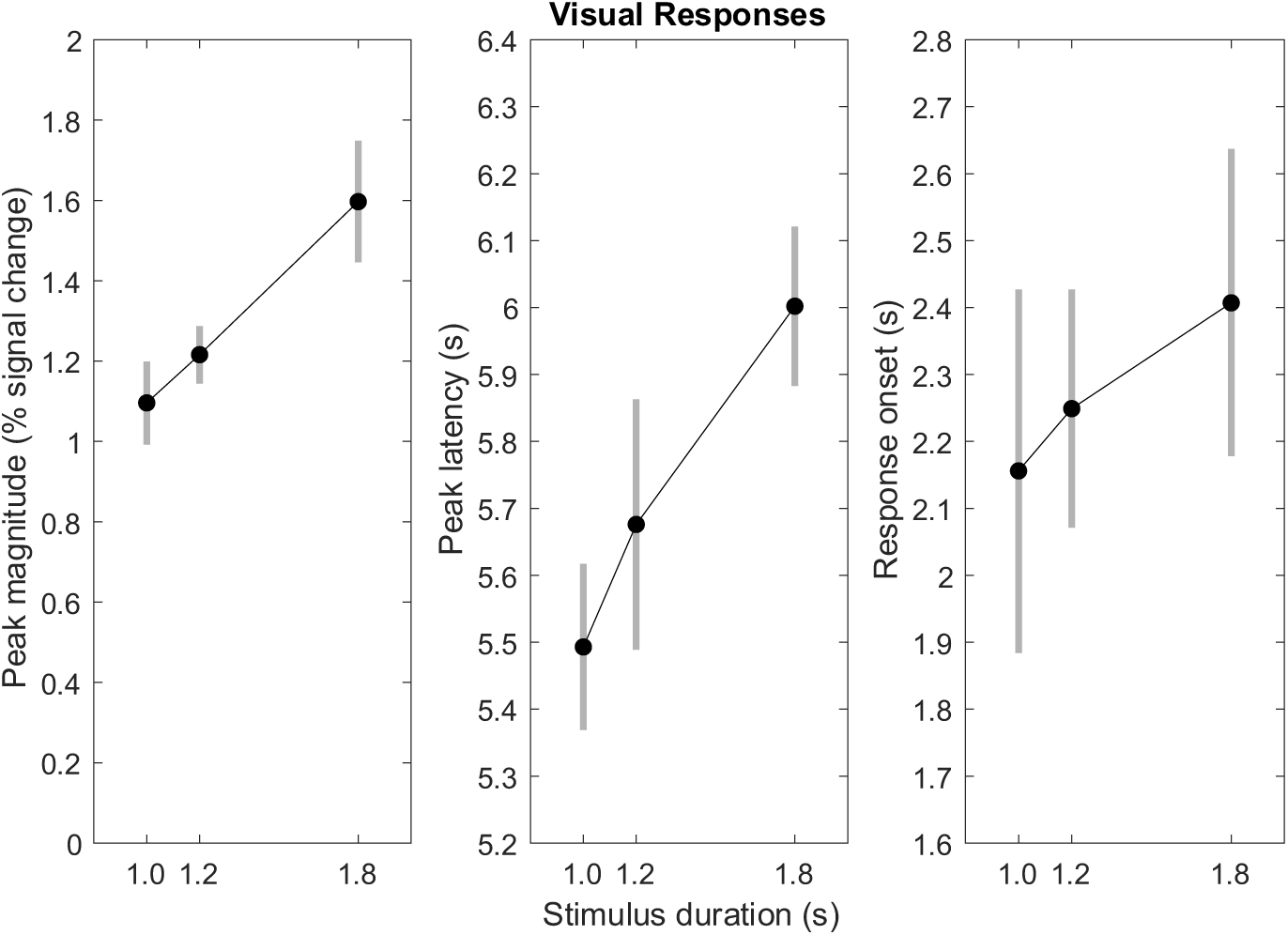
Estimated HRF parameters from visual stimulation across different durations. 600ms (1200ms-1800ms) and 800ms (1000ms-1800ms) increases in visual stimulus durations led to significant increases in the mean peak magnitude (left column) and peak latency (middle column) of their corresponding fitted hemodynamic responses. There were negligible effects of stimulus duration on response onset (right column). Error bars represent 95% confidence intervals of the mean.

Critically, increases in visual stimulus durations also led to significant increases in peak latency. Peak latency increased by 0.18 s (SD = 0.26, 95% CI [0.03 0.33]) as stimulus duration increased by 200 ms (from 1000 to 1200 ms), and it increased by 0.32 s (SD = 0.49, 95% CI [0.04 0.60]) as stimulus duration increased by 600 ms (from 1200 to 1800 ms). The overall peak latency increase of 0.50 s (SD = 0.18, 95% CI [0.40 0.61]) was similar to the increase observed in Experiment 1; the key difference was the much lower variance in Experiment 2.

In contrast, increases in stimulus intensity affected only the HRF’s peak magnitude, not its peak latency (Figure 11, top right; Figure 13). The increase in stimulus contrast from 10% to 100% produced an 0.22% (SD = 0.18, 95% CI [0.12 0.32]) increase in peak magnitude. This same stimulus change, however, produced a non-significant change of −0.20s (SD = 0.49, 95% CI [-0.48 0.07]) in peak latency. Parameter estimates and their variances for each condition combination can be found in the Supplementary Materials as csv tables.

**Figure 13.**
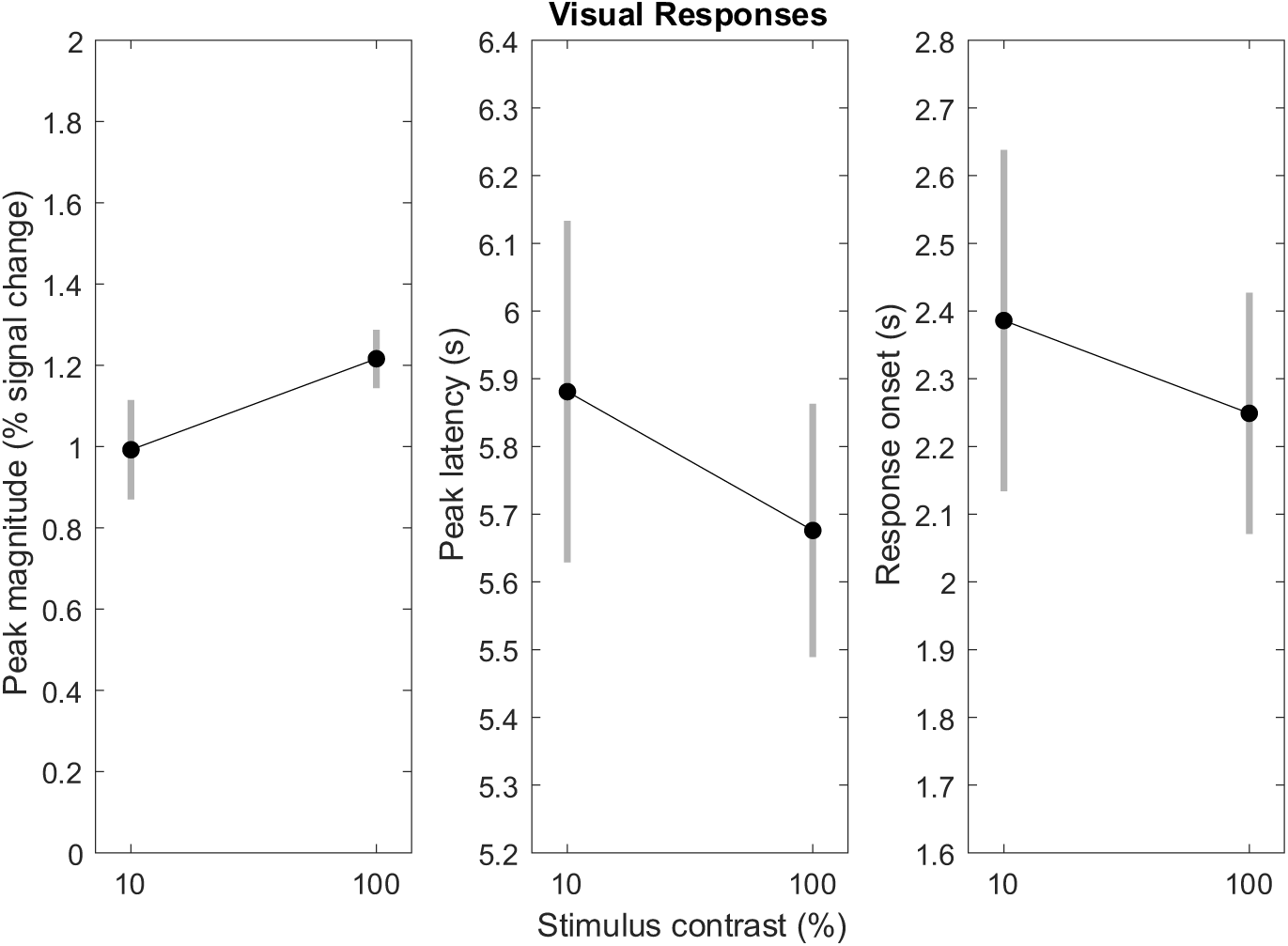
Estimated HRF parameters from visual stimulation across different intensities. Increasing visual stimulus intensity (contrast) led to a significant increase in the mean peak magnitude (left column) of the corresponding fitted hemodynamic responses. There were negligible effects of stimulus intensity on peak latency (middle column) and response onset (right column). Error bars represent 95% confidence intervals of the mean.

#### 3.2.2. Peak latency differentiates auditory stimulus duration and intensity manipulations

Increases in auditory stimulus durations led to significant increases in both peak magnitudes and peak latencies of the hemodynamic response (Figure 11, bottom left; Figure 14). Peak magnitude increased by 0.09 (SD = 0.16, 95% CI [0.004 0.19]) as stimulus duration increased by 200 ms (from 1000 to 1200 ms), and it increased by 0.18 (SD = 0.24, 95% CI [0.04 0.31]) as stimulus duration increased by 600 ms (from 1200 to 1800 ms). The overall peak magnitude increase of 0.27 (SD = 0.09, 95% CI [0.22 0.33]) was similar to the increase observed in Experiment 1 for the same stimulus duration change, and both were smaller than their visual counterparts.

**Figure 14.**
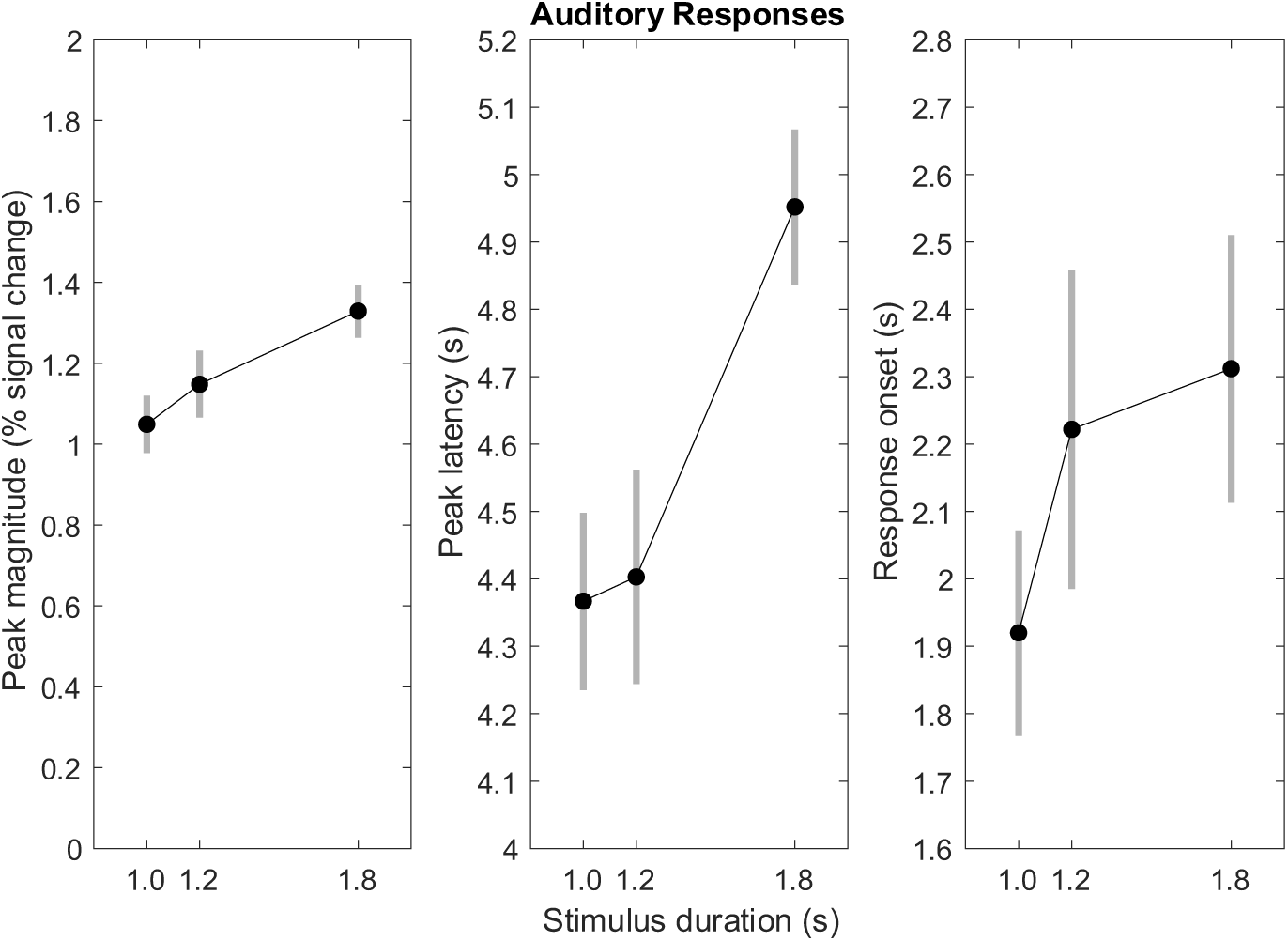
Estimated HRF parameters from auditory stimulation across different durations. 600ms (1200ms-1800ms) and 800ms (1000ms-1800ms) increases in auditory stimulus durations led to significant increases in the mean peak magnitude (left column) and peak latency (middle column) of their corresponding fitted hemodynamic responses. There were negligible effects of stimulus duration on response onset (right column). Error bars represent 95% confidence intervals of the mean.

Critically, increases in auditory stimulus durations also led to increases in peak latency, which were statistically significant for the 600 ms duration change. Peak latency increased by 0.03 s (SD = 0.33, 95% CI [-0.15 0.22], non-significant) as stimulus duration increased by 200 ms (from 1000 to 1200 ms), and it increased by 0.54 s, (SD = 0.48, 95% CI [0.27 0.82]) as stimulus duration increased by 600 ms (from 1200 to 1800 ms). The overall peak latency increase of 0.58s (SD = 0.15, 95% CI [0.49 0.67]) was considerably larger and less variable than the increase observed in Experiment 1. It was comparable, however, to the peak latency increases observed for visual stimuli.

Increases in stimulus intensity had no significant effects on either the HRF’s peak magnitude or its peak latency (Figure 11, bottom right; Figure 15). The increase in stimulus volume from 30% to 90% of the speaker’s maximum produced an 0.07% (SD = 0.18, 95% CI [-0.02 0.18]) increase in peak magnitude. This same stimulus change also produced a change of −0.11s (SD = 0.26, 95% CI [-0.26 0.03]) in peak latency.

**Figure 15.**
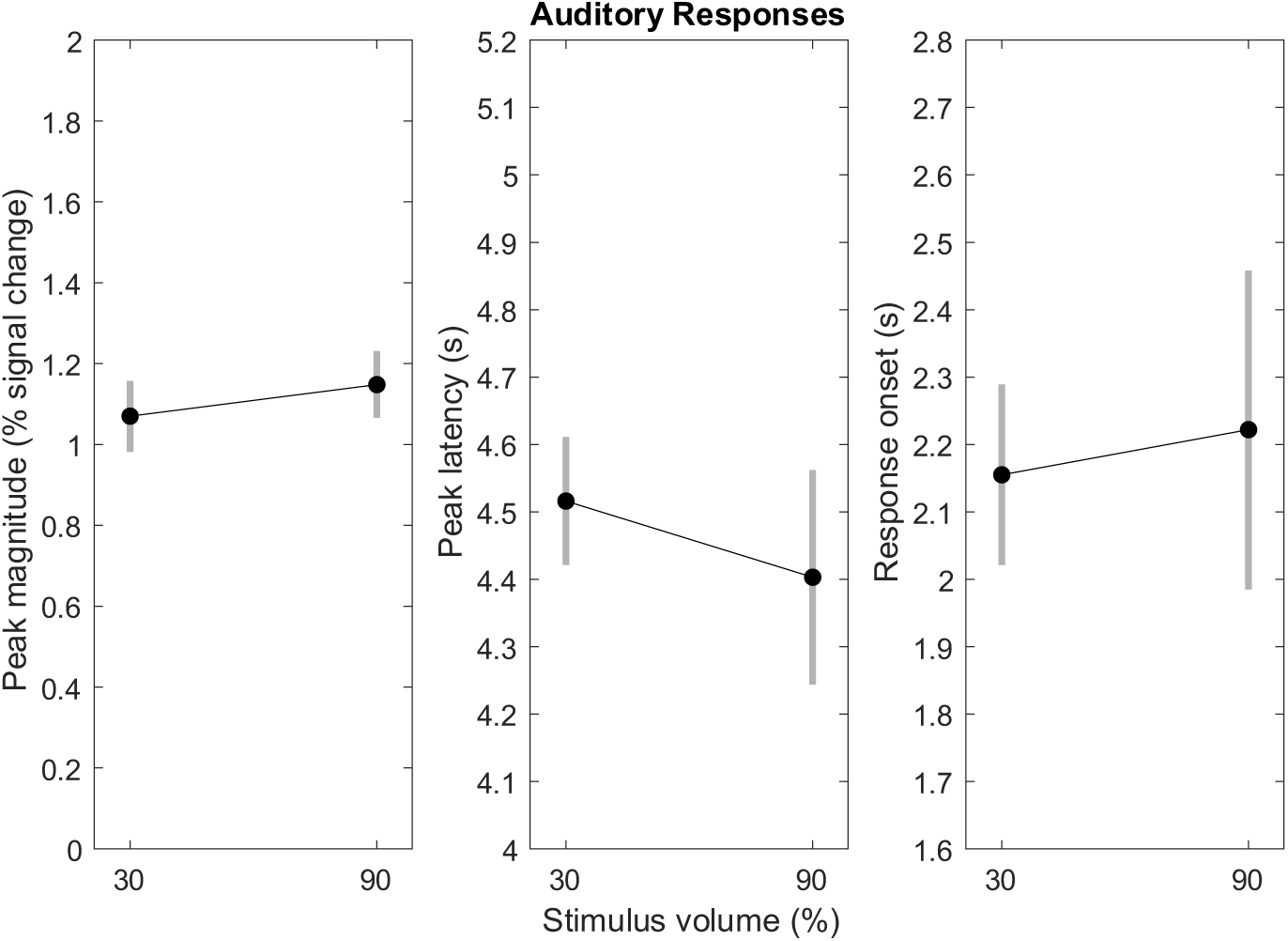
Estimated HRF parameters from auditory stimulation across different intensities. Increasing auditory stimulus intensity (volume) led to negligible effects on the peak magnitude (left column), peak latency (middle column) and response onset (right column) of the corresponding fitted hemodynamic responses. Error bars represent 95% confidence intervals of the mean.

## 4. General Discussion

### 4.1. Summary of Findings

We predicted that increases in stimulus duration would affect both the peak magnitude and peak latency of the evoked hemodynamic response. In contrast, increases in stimulus intensity would affect only peak magnitude. The results of Experiments 1 and 2 support these predictions, demonstrating that event-related BOLD responses can reliably index sub-second differences in stimulus duration when the underlying signal is sufficiently robust.

In Experiments 1 and 2, we explored whether the BOLD responses to brief visual and auditory stimuli could be distinguished using peak magnitude and peak latency parameters from fitted HRFs. In Experiment 1, significant differences in peak magnitude were observed across stimulus durations when stimulus intensity was high, suggesting that even a 200 ms duration difference could be detectable. However, signal variability and low response amplitudes made drawing timing-related conclusions difficult. In Experiment 2, we improved signal quality by increasing stimulus durations (1000, 1200, and 1800 ms), reducing the number of conditions, and using a longer TR. These changes yielded clearer effects in both modalities: Peak latency and peak magnitude showed robust effects for 600 ms stimulus duration differences, with some effects significant even for 200 ms differences. Compared to Experiment 1, similar stimulus conditions in Experiment 2 (e.g., 900-1000 ms stimulus durations) yielded more stable parameter estimates and narrower confidence intervals, likely reflecting the combined effects of improved task and scanning parameters.

Our findings held across modalities, suggesting a generalizable dissociation between the effects of intensity and duration on the BOLD response to brief stimuli. That said, the timing effects with visual stimuli were generally more robust, showing more consistent peak latency shifts and narrower confidence intervals across conditions. This asymmetry may reflect both environmental and intrinsic factors. The scanner environment poses greater challenges for auditory stimulation, with background noise and imperfect stimulus delivery reducing the reliability of evoked responses. By contrast, visual cortex is known to produce robust BOLD responses to simple stimuli (Boynton et al., 2012; Goodyear & Menon, 1998), likely due to its clear retinotopic organization and strong hemodynamic coupling. Although our study was not designed to test modality differences directly, future work may benefit from optimizing stimulus presentation and analysis for specific brain regions of interest, including auditory cortex.

### 4.2. BOLD timing reflects stimulus duration, validating fMRI for mental chronometry

Our findings support existing hemodynamic models by demonstrating that changes in stimulus intensity and duration exert dissociable effects on the BOLD signal (Boynton et al., 2012; Grinband et al., 2008). They also extend the applicability of these models to brief stimulation periods (< 2000 ms) and sub-second timing differences (as small as 200 ms).

Changes in stimulus intensity induced a non-linear scaling of peak magnitude without affecting peak latency, whereas changes in stimulus duration affected both peak magnitude and peak latency simultaneously. These patterns held across experiments and sensory modalities, with consistent parameter estimates across trials and participants—suggesting that sub-second BOLD timing shifts are detectable on a timescale meaningful for cognitive events (Chen et al., 2023; Grinband et al., 2008; Miezin et al., 2000).

This dissociation supports the use of fMRI for mental chronometry. By showing that BOLD peak latency shifts track stimulus duration but not intensity, we strengthen the foundation for inferences about processing time based on peak timing. Several prior studies have adopted this logic to identify cognitive bottlenecks (Asplund et al., 2010; Dux et al., 2006, 2009; Marti et al., 2012; Scalf et al., 2011; Tombu et al., 2011; Vagharchakian et al., 2012), including recent multitasking work with ultrafast fMRI (Yue et al., 2025). Our results reinforce the interpretability of peak latency differences and provide empirical support for a widely used, but previously unvalidated, assumption in time-resolved fMRI.

Beyond validating peak latency as a marker of processing duration, our findings align with recent advances in extracting fast neural dynamics from the hemodynamic signal. Improvements in acquisition and analysis have enabled increasingly fine-grained temporal measurements using fMRI, including Granger causality-based lag analyses (Katwal et al., 2013; Rogers et al., 2010; Wang et al., 2017), neural sequence decoding (Li et al., 2024; Setzer et al., 2022; Wittkuhn & Schuck, 2021; Yue et al., 2025), and model-based comparisons of dynamic processes (Marxen et al., 2023). High-field and high-sampling-rate fMRI methods (Chen et al., 2023; Dowdle et al., 2021; Lewis et al., 2018; Polimeni & Lewis, 2021) have further improved signal quality and temporal resolution, allowing for greater precision in characterizing brief neural events. While many of these approaches focus on tracking response onsets or sequences of activation, our findings broadly clarify the conditions under which peak latency shifts reflect changes in neural duration rather than intensity. This distinction enhances the interpretability of latency-based measures and contributes to the conceptual foundation of temporally-sensitive fMRI techniques—including functional connectivity analyses (Smith et al., 2011) and dynamic causal modelling (Stephan et al., 2007).

### 4.3. Parameterized HRFs, bootstrap resampling, and good design improve inference

Hemodynamic responses to brief stimuli often yield noisy data, particularly when stimulus intensity is low. To improve robustness and extract interpretable timing metrics, we fitted a parameterized HRF to the event-related signals for each participant by condition. Each HRF included the timing-sensitive parameters of onset, delay, and dispersion. These parameters are not orthogonal and so can trade-off; they also do not uniquely relate to neural duration (Lindquist et al., 2009; Wager et al., 2005). As such, we computed peak latency directly from the fitted response curve, an approach that produced interpretable and robust estimates. More generally, the success of this approach in capturing reliable latency differences supports the use of variable-epoch models in fMRI analyses (Grinband et al., 2008), which allow both timing and amplitude to vary across conditions (Lindquist et al., 2009). By contrast, variable-impulse models treat only the amplitude of the BOLD response as flexible.

Many prior studies using brief stimuli have focused on HRF amplitude instead of timing (Yeşilyurt et al., 2008), and some have failed to find significant differences in peak latency even when they might be expected (Lewis et al., 2018). These limitations may reflect shortcomings in conventional HRF modelling, especially for brief events on sub-second timescales (Polimeni & Lewis, 2021). In contrast, the approach adopted here—fitting a parameterized HRF to event-related BOLD signals using nonlinear least squares—recovered consistent timing effects from stimulus duration differences as small as 200 ms. The method is simple, efficient, and easily interpretable: It tests how well the observed task-evoked BOLD signal conforms to an *a priori* response function, without requiring additional model complexity or custom basis sets.

Although fitting parameterized HRFs improved robustness, variability in the timecourses still led to unstable parameter estimates—especially for timing metrics like peak latency. In Experiment 1, this problem was acute: Hemodynamic responses for many participant-by-condition combinations lacked a clear peak, and the resulting peak latency estimates were highly variable and often unreasonable. These noisy estimates exerted an outsized influence on the summary statistics. To address this issue, we used bootstrap resampling to estimate parameter means and variances, reasoning that this approach would mitigate outlier influence and enhance inferential reliability.

Bootstrap resampling improved the reliability of the parameter estimates, revealing effects that standard approaches missed. For example, at 10% visual stimulus contrast, parametric analyses found no difference in peak latency between 300 ms and 900 ms stimuli—both estimates were 5.48 s—but the standard error was larger for the low-signal 300 ms condition (0.64 s versus 0.26 s). Bootstrapped estimates, by contrast, showed a reliable 0.45 s difference, with smaller and more consistent standard errors (0.33 s and 0.20 s).

Estimate precision improved as well. Across all auditory conditions, bootstrapped confidence intervals for auditory peak latency estimates were 26% narrower than their parametric counterparts on average; for peak latency differences, the reduction was 43%. This improved precision translated into clearer inference. In Experiment 1’s 100% contrast condition, the estimated peak latency difference between 100 ms and 900 ms visual stimuli was 0.62 s (95% CIs: [0.06 1.17]) using parametric methods, and 0.72 s (95% CIs: [0.30 1.18]) using bootstrapping—a notable improvement in inferential clarity.

While bootstrap resampling improved parameter stability in Experiment 1, it could not fully overcome the limitations imposed by weak or noisy evoked responses. The design enhancements in Experiment 2—longer stimuli, fewer conditions, and a longer TR— produced stronger evoked signals and more reliable estimates, largely obviating the need for resampling. Improvements to experimental design and signal quality tend to yield greater returns than post hoc analytic corrections and should be prioritized when feasible.

Nonetheless, resampling remains a valuable and easily implemented complement, especially when studying subtle timing effects near the resolution limits of fMRI.

## 5. Conclusion

Our study empirically demonstrated that the predicted effects of stimulus duration and intensity on evoked hemodynamic responses can be dissociated, even for sub-second duration changes. For both visual and auditory stimuli, duration changes affected both peak latency and peak magnitude, whereas intensity changes affected peak magnitude alone. When the underlying hemodynamic responses are robust, duration changes as small as 200 ms can produce significant changes in peak latency.

These findings support the validity of existing hemodynamic models and extend their applicability to brief stimuli and fine-grained timing differences—conditions common in cognitive neuroscience. Careful experimental design and improved HRF modelling may further enhance sensitivity in such contexts. Although we used perceptual manipulations in sensory cortices, we expect that cognitive manipulations would yield similar effects on event-related hemodynamic responses in other brain regions. By clarifying and confirming the conditions under which BOLD timing reflects stimulus duration, our study provides a principled foundation—and an accessible approach—for using time-resolved fMRI as a tool for mental chronometry.

## Data and code availability

Means and variances for peak amplitude, peak latency, and onset latency for each condition combination is provided in csv files with this submission. All data and code is currently available upon reasonable request, and we will make such information, as well as other parameter estimates, publicly available before publication.

## Author Contributions (CRediT)

Alvin P.H. Wong: Conceptualization, Methodology, Visualization, Formal analysis, Writing – Original Draft, Writing – Review & Editing.

Esther X.W. Wu: Data curation, Investigation, Visualization, Writing – Review & Editing. Baxter P. Rogers: Methodology, Resources, Writing – Review & Editing.

Christopher L. Asplund: Conceptualization, Methodology, Supervision, Funding acquisition, Writing – Original Draft, Writing – Review & Editing.

## Supporting information

Expt 1 parameter means and variances (auditory)

Expt 1 parameter means and variances (visual)

Expt 2 parameter means and variances

## Acknowledgements

The authors would like to thank Stuart W.G. Derbyshire, John J. Totman, Stevia Ng Gogna, Mary C. Stephenson, and Paul E. Dux for helpful discussions.

